# The cell polarity landscape in Arabidopsis male and female gametophytes

**DOI:** 10.64898/2026.10.30.763139

**Authors:** Samuel Koh Wee Han, Monique van Schie, Catherine Albrecht, João Jacob Ramalho, Bodhi Voss, Dolf Weijers

## Abstract

Plant cell polarity underlies the capacity to directionally transport auxin and other small molecules, but also precedes asymmetric cell division in the stomatal lineage. Except for specific contexts, mechanisms underlying cell polarization are poorly understood. Most insights on plant cell polarity have come from studies on flowering plant sporophytes, while no polar protein localization has yet been reported in flowering plant gametophytes, despite clear anisotropic growth and asymmetric cell divisions in this generation. Inspired by the identification of deeply conserved SOSEKI proteins and their associated partners that mark polar domains in bryophyte gametophytes and Arabidopsis sporophytes, we here investigate cell polarity in Arabidopsis gametophytes. We discover a rich landscape of polar domains throughout male and female gametophyte development, marked by one or more polar proteins. We find that successive steps in male gametophyte polarization can be mechanistically uncoupled, and the first polarization step precedes asymmetric cell division. Using ectopic expression, we find that there is a near-universal capacity for polar protein localization in gametophytes through a single polarity system. Our work offers the first insight into gametophyte cell polarity in Arabidopsis, identifies novel polar domains, and presents a starting point for future dissection of mechanisms underlying plant cell polarity.

## Introduction

Cell polarity helps spatially organize cells. While in single-celled organisms, this can aid in cell division, migration or local differentiation (e.g. flagella) (Chiou et al., 2017; Shapiro et al., 2009; Treuner-Lange and Søgaard-Andersen, 2014), in multicellular organisms, cell polarity aids coordinated cell behavior and morphogenesis. Polar domains can create cell adhesions, organize directional transport and enable oriented and asymmetric cell division (Knoblich, 2010; Rodriguez-Boulan and Macara, 2014). Polarity at the cellular level is marked by the local accumulation of proteins, RNA, organelles, extracellular matrix or membrane components (Martin and Ephrussi, 2009; St Johnston and Ahringer, 2010). Mechanisms guiding such local accumulation therefore organize the polarity system. Mechanisms underlying cell polarization in animals and yeasts have been studied in some detail and have led to the identification of key pathways (Chiou et al., 2017; Nelson, 2003; St Johnston and Ahringer, 2010). Several polarity-localized proteins have been identified in plants, and these include the PIN-FORMED auxin transporters (Gälweiler et al., 1998; Wisniewska et al., 2006), BOR1 and NIP5 borate transporters (Takano et al., 2006; Takano et al., 2010; Wang et al., 2017), Rho of Plants (ROP) GTPases (Fu et al., 2005; Xu et al., 2010), as well as BASL, POLAR and BRX scaffold proteins (Dong et al., 2009; Houbaert et al., 2018; Koh et al., 2021; Marhava et al., 2018). Essentially all of these plant polarly localized proteins are plant-specific and do not share clear homology with non-plant polar proteins (Dettmer and Friml, 2011; Ramalho et al., 2022). Likewise, the regulators of animal and yeast cell polarity are missing from plant genomes (Dettmer and Friml, 2011; Yang, 2008). Thus, plants must have invented their own cell polarity system, and its identification offers an interesting challenge. More recently, SOSEKI proteins were identified as edge-polarized proteins, conserved across land plants, and sharing homology with animal Dishevelled polarity proteins (van Dop et al., 2020; Yoshida et al., 2019). Whether this is a rare case of convergent evolution, or if there are more unexplored homologies between animal and plant polarity systems, is an open question.

Almost all studies on plant cell polarity were performed in the flowering plant Arabidopsis, or more generally in vascular plants (Dettmer and Friml, 2011; Ramalho et al., 2022; Yang, 2008). Land plants are remarkable in that they have a clear alternation between generations: a haploid gametophyte generation and a diploid sporophyte generation. The earliest land plant gave rise to two domains: non-vascular plants (Bryophytes) and vascular plants (Tracheophytes). The former group includes mosses, liverworts and hornworts, the latter group includes lycophytes ferns, seed plants and flowering plants (Harris et al., 2022; One Thousand Plant Transcriptomes Initiative, 2019; Puttick et al., 2018). A defining distinction between the two groups is that in bryophytes, the haploid gametophyte is dominant and indeterminately growing, and the sporophyte is reduced to a diminutive structure. Inversely, with ferns being the only exception, in tracheophytes, the sporophyte dominates and grows indeterminately, while the gametophyte is reduced to a structure of but a few cells. As mentioned, cell polarity has mostly been studied in flowering plants, and exclusively in sporophytes, with the exception of tip-growing pollen tubes, which reflect a unique type of polarity that may not reflect interphase polarity in non tip-growing cells (Hepler et al., 2001; Qin and Yang, 2011). It is therefore an open question whether the tracheophyte sporophyte, which has complex tissue arrangements, has unique need for cell polarity systems. The recent finding that SOSEKI proteins mark polar domains in the Bryophyte gametophyte (Pukhovaya et al., 2026; van Dop et al., 2020), suggests that sporophytic and gametophytic cell polarity have a common origin. The proteomic identification of common, polarized SOSEKI-associated proteins in the sporophyte of the flowering plant *Arabidopsis thaliana* and the gametophyte of the liverwort *Marchantia polymorpha* (Pukhovaya et al., 2026) further supports the notion of a deeply conserved polarity mechanisms across the two generations of all land plants. However, despite decades of research on flowering plant gametophytes, there are no reports on cell polarity during their development.

We here explore polarity in the male and female gametophytes of the flowering plant *Arabidopsis thaliana*. Both are multicellular: male gametophytes form from microspores, following male meiosis (Berger and Twell, 2011; Borg et al., 2009; Twell, 2011). They undergo a first, asymmetric division to generate a vegetative cell and a generative cell. The latter divides once more to generate two sperm (Berger and Twell, 2011; Twell et al., 1998). The female gametophyte forms from a megaspore, the survivor of the 4 meiotic products (Drews and Koltunow, 2011; Schmidt et al., 2015; Yadegari and Drews, 2004). Following anisotropic growth, it divides 3 times to generate 8 nuclei that are patterned in an embryo sac which, upon cellularization, contains two synergids, an egg cell, a central cell and three antipodal cells. The structure is highly polarized, but has an unusual developmental progression that includes a coenocytic stage before cellularization (Drews and Koltunow, 2011; Sprunck and Gross-Hardt, 2011). Thus, both male and female gametophyte feature polarized structure, as well as asymmetric cell division and/or polar patterning. However, whether this brief phase of flowering plant development is marked by cell polarity remains an entirely open question. Microspores are isolated from their sporophytic maternal tissue, the tapetum (Scott et al., 2004), while female gametophytes grow within sporophytic maternal tissue, the nucellus (Ceccato et al., 2013; Pagnussat et al., 2009), in which polar PIN protein localization has been reported. As mentioned, it is however, an open question whether the gametophytes themselves are polarized.

Starting from a large collection of SOSEKI proteins and SOSEKI-associated proteins, we here report that both male and female gametophytes are profoundly polarized. Both contain multiple polar domains, marked by different proteins. In the male, we show that polar SOSEKI protein localization precedes the earliest known sign of symmetry breaking. We show that successive polarization steps can be mechanistically uncoupled. Lastly, we show that a single polarity system appears to guide the localization of all proteins studied. This work reports a rich landscape of cell polarization in the haploid generation in Arabidopsis and will mark a starting point for the dissection of mechanisms guiding cell polarity in plants.

## Results

### SOSEKI proteins mark polar domains during female reproductive development

We initially explored the expression domains and subcellular localization patterns of the Arabidopsis SOK family protein during female gametophyte development, and in the sporophytic tissues in which the gametophyte is enclosed. For both this analysis and the analysis presented in the following sections, localization patterns are summarized in Figure 8.

During stage I of female gametogenesis, the base of the ovule is formed by the formation of finger-like protrusions of the placenta (Robinson-Beers et al., 1992; Schneitz et al., 1995). At this stage, SOK2, 3, and 5 are expressed, each polarized inwards to the center of the developing ovule (Fig. 1F, K, U). During stage II, the centrally localized Megaspore Mother Cell (MMC) will undergo meiosis to form the tetrad. At the same time, the inner and outer integuments are initiated from sporophytic cells (Robinson-Beers et al., 1992; Schneitz et al.,

**Figure 1.**
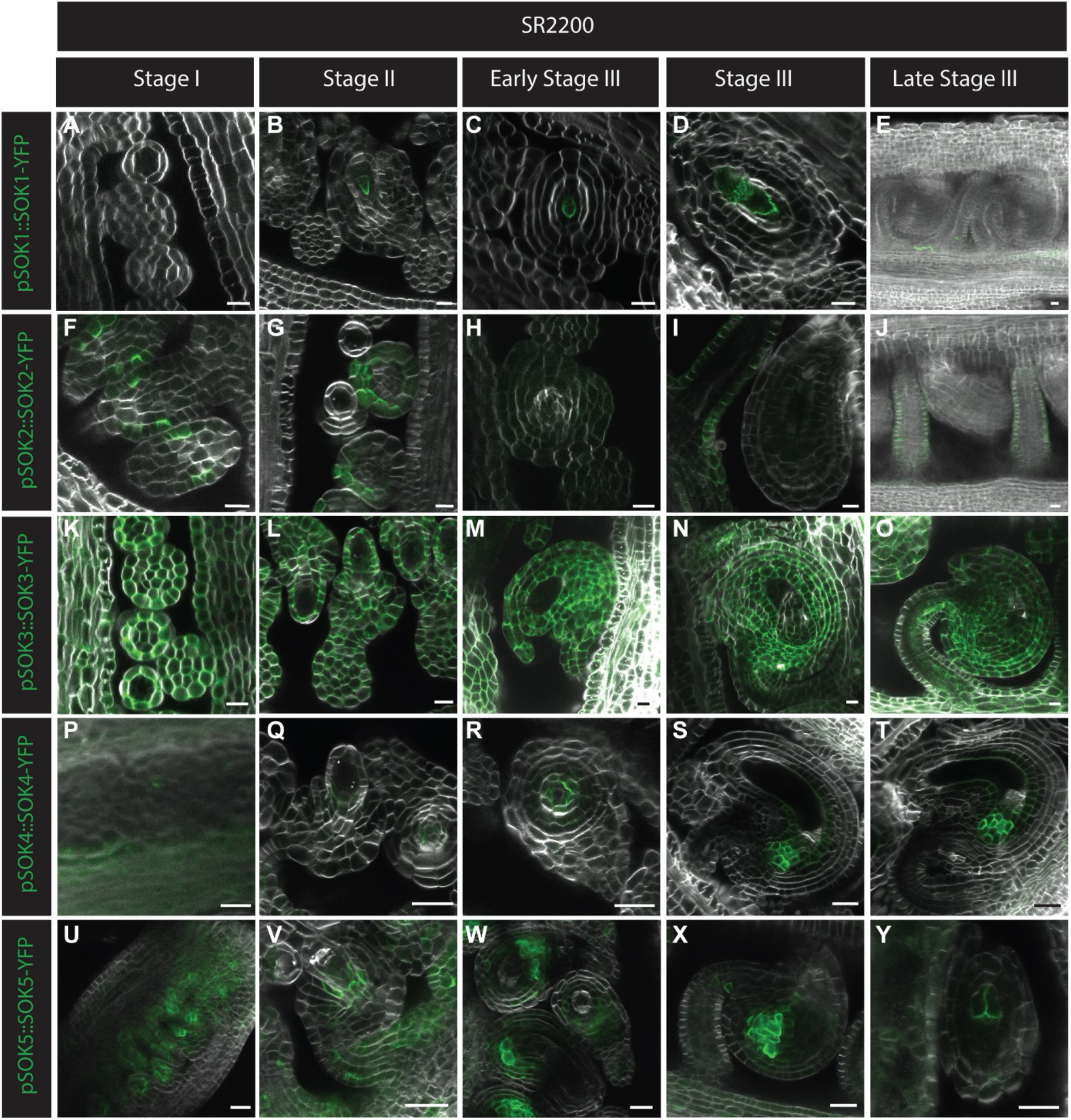
All SOK proteins are expressed and polarly localized during ovule development, only SOK1 is restricted to the female gametophyte. (A-Y) Confocal sections of the developing ovule in stage I, stage II and III expressing pSOK1::gSOK1-YFP (A-E), pSOK2::gSOK2-YFP (F-J), pSOK3::gSOK3-YFP (K-O), pSOK4::gSOK4-YFP (P-T) and pSOK5::gSOK5-YFP (U-Y). YFP, green; SR-2200, white. Images are displayed with brightness normalization and in some cases rotated or inverted to facilitate comparison of images. Tissues were fixed, cleared using Clearsee alpha (Kurihara et al., 2021) and stained with SR-2200 dye (Musielak et al., 2015) before imaging. Scale bars, 10 µm.

1995). We found all five SOKs to be expressed around stage II. Since we did not explicitly address division of the MMC, we can not distinguish pre-versus post-meiotic patterns. Only SOK1 is expressed in the MMC (or megaspore), while the other 4 proteins mark sporophytic cells (Fig. 1, S1). SOK1 is polarly localized towards the chalazal pole of the MMC/megaspore (Fig. 1B). SOK4 and 5 are both expressed in the cells neighboring the MMC/megaspore and polarized towards it, with SOK5 in a broader expression pattern (Fig. 1Q, V). SOK3 is ubiquitously expressed but seems polarized towards the MMC/megaspore in the neighboring cells (Fig. 1L). SOK2 marks the outer edge cell layers of the funiculus (Fig. 1G).

During early stage III, the tetrad degenerates and only one nucleus remains in the newly formed embryo sac. The outer integument starts to envelop the the structures underneath it. The nucleus undergoes three rounds of mitosis while the surrounding tissues grow and start to curve the ovule by differential growth rates. In late stage III, the eight nuclei in the embryo sac cellularize to form the the egg cell, antipodals, synergids and central cell. Lastly, two nuclei within the central cell fuse and the antipodal cells degenerate (Robinson-Beers et al., 1992; Schneitz et al., 1995). During early stage III, SOK1 is expressed in the embryo sac, polar towards the chalazal end, while at the last part of stage III the signal fades (Fig. 1C-E). During most of stage III, SOK4 and 5 are expressed in the nucellus, polarized towards the embryo sac (Fig. 1R-T, W-X). SOK2 keeps marking the outer edge cells of the funiculus (Fig. 1I, J), while SOK3 is still broadly expressed (Fig. 1M-O). Following cellularization of the embryo sac, we find only SOK5 to be expressed, and polarized inward in the two synergids (Fig. 1Y).

Thus, Arabidopsis female gametophyte development is marked by a landscape of polar domains, both within the developing coenocytic gametophyte and the cellularized stage, and in the surrounding sporophytic tissue.

### SOSEKI proteins mark symmetry breaking and polar domains during male gametophyte development

Microspores undergo symmetry breaking, followed by nuclear migration (Oh et al., 2016; Park et al., 1998) and pollen mitosis I (PMI), which divides the polarized microspore asymmetrically into a large vegetative cell and a small generative cell (Eady et al., 1995; Mizuta et al., 2026). The generative cell divides again at pollen mitosis II (PMII) to form two sperm cells (Berger and Twell, 2011; Twell et al., 1998).

To characterize the developmental distribution of SOK proteins during male gametophyte development, we imaged YFP fusions of all five Arabidopsis SOK proteins expressed under their native promoters. Pollen was counterstained with Hoechst 33342 to identify nuclei and assign the developmental stage (Fig. 2).

**Figure 2.**
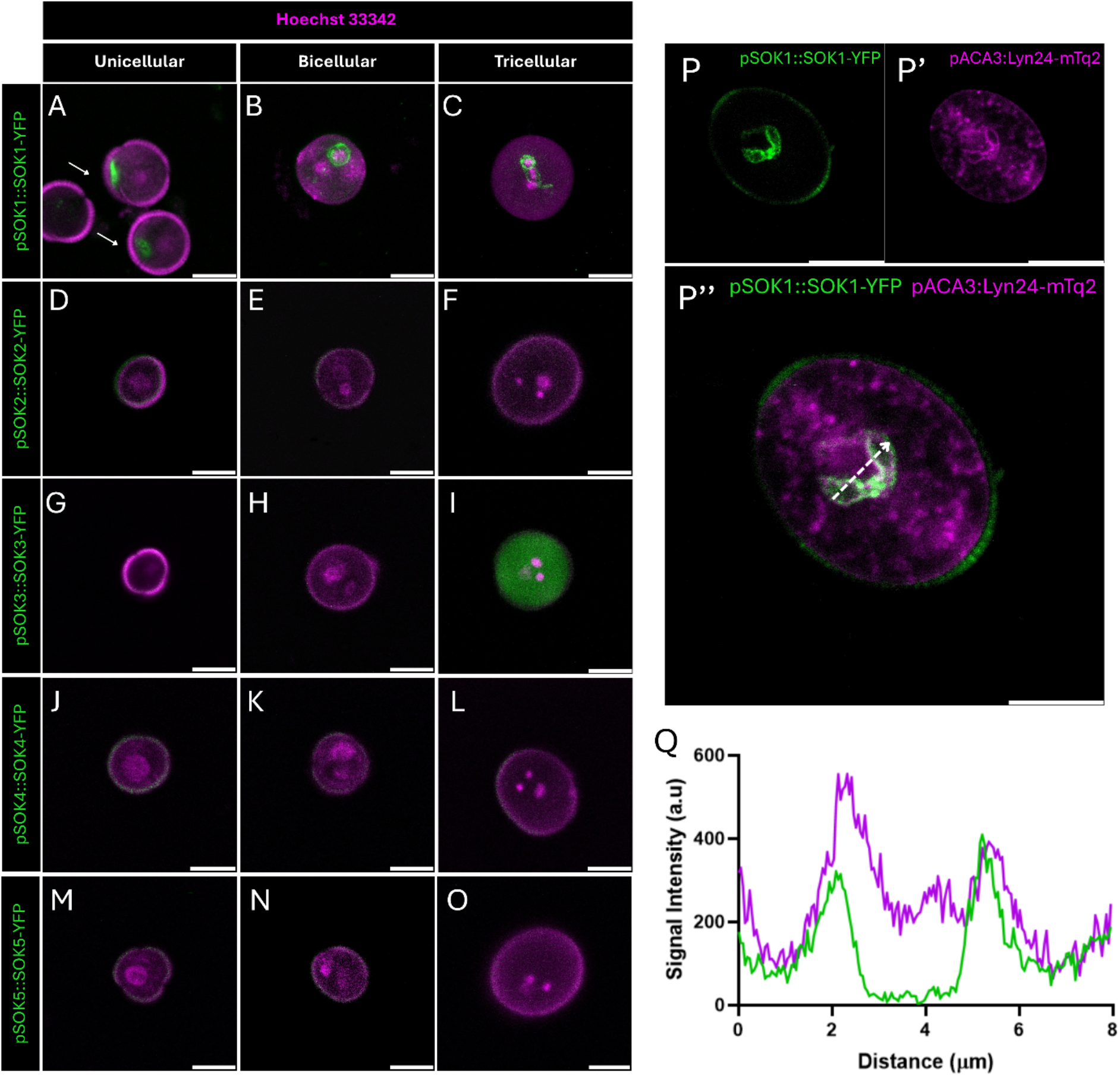
SOK1 is the only SOSEKI family member with a polar, membrane-associated distribution in the male gametophyte. (A–O) Confocal sections of unicellular microspores, bicellular pollen and tricellular pollen expressing pSOK1::gSOK1-YFP (A–C), pSOK2::gSOK2-YFP (D–F), pSOK3::gSOK3-YFP (G–I), pSOK4::gSOK4-YFP (J–L) and pSOK5::gSOK5-YFP (M–O). YFP, green; Hoechst 33342, magenta. Arrows mark the polar cortical domain of SOK1-YFP. (P–P’’) SOK1-YFP with the peri-germ cell membrane marker pACA3::Lyn24-mTurquoise2 in tricellular pollen. (Q) Fluorescence intensity profile along the line indicated in P’’. Images were acquired under identical settings and are displayed with brightness normalization. Scale bars, 10 µm.

Among the five SOK proteins, only SOK1 and SOK3 were detected during pollen development (Fig. 2). SOK1 showed a stage-dependent pattern. In unicellular microspores, it accumulated in a discrete crescent at one side of the cell periphery before PMI (Fig. 2A). After PMI, it outlined the generative cell (Fig. 2B). After PMII, it surrounded the two sperm cells (Fig. 2C). SOK3-YFP was undetectable in unicellular and bicellular pollen but gave a diffuse cytosolic signal in the vegetative cell of tricellular pollen (Fig. 2G–I). After PMI, the generative cell is internalized by the vegetative cell. The generative cell, and later the sperm cells, are therefore enclosed by a membrane derived from the vegetative cell plasma membrane. This membrane is called the peri-germ cell membrane (PGCM), or inner vegetative plasma membrane (Sugi et al., 2023; Sugi et al., 2024). The SOK1 pattern suggests that it may localize to this PGCM. To test this, we combined SOK1-YFP with the PGCM marker pACA3::Lyn24-mTurquoise2 (Motomura et al., 2021; Sugi et al., 2023). SOK1-YFP closely overlapped with Lyn24-mTurquoise2 around the germline compartment (Fig. 2P), and fluorescence intensity profiles across the labelled membrane showed peaks of both reporters at the same positions (Fig. 2Q). These observations are consistent with SOK1 localization to the PGCM during post-PMI development. However, given the close proximity of the PGCM to the germ cell plasma membrane (Sugi et al., 2023), we can not exclude that some of the SOK1 signal is derived from the germ cell plasma membrane.

We conclude that SOK1 marks an early step in symmetry breaking in the microspore, prior to nuclear migration, and that it marks all successive steps in engulfment of the germ cell and sperm, thus reporting on several key events associated with male gametophyte polarization.

### Functional diversification of SOSEKI proteins reveals two-step polarization in the male gametophyte

Among the five Arabidopsis SOK proteins, only SOK1 polarizes in the male gametophyte, where it marks two very different steps: symmetry breaking in the unpolarized microspore and engulfment of the generative cell. It is unclear whether these two steps are part of the same, continuous process, and if SOK1 protein is unique, or whether it reflects a generic SOK property. To test the latter, we expressed each of the other 4 SOK proteins as YFP fusions from the SOK1 promoter, alongside SOK1 as a control. We introduced these into the nuclear FB037 reporter line, which contains markers for nuclei (pACT11::H2B-mRFP) and germline nuclei (pMGH3::MGH3-eGFP).

At the polarization stage, the late unicellular stage before PMI, SOK1 accumulated in a polar domain at the cell periphery (Fig. 3A–B). Strikingly, SOK2, SOK3, SOK4, and SOK5 could all localize to this same domain when expressed in the young microspore (Fig. 3F, J, N, R). Thus, the paralogs can adopt an early polar distribution resembling that of SOK1. After PMI, all paralogs diverged from SOK1. While SOK1 surrounded the generative cell in bicellular pollen (Fig. 3C) and the sperm cells in tricellular pollen (Fig. 3D), SOK2 (Fig. 3G), SOK3 (Fig. 3K), SOK4 (Fig. 3O) and SOK5 (Fig. 3S) were mainly cytosolic and lacked the membrane enrichment that SOK1 shows at the PGCM. In tricellular pollen, we detected no signal from any of the SOK1 paralogs in the germline. Thus, when expressed in the SOK1 domain, the paralogs can form an early polar pattern but not localize to the PGCM. This identifies differences among SOK proteins but also shows that the two steps are discontinuous.

**Figure 3.**
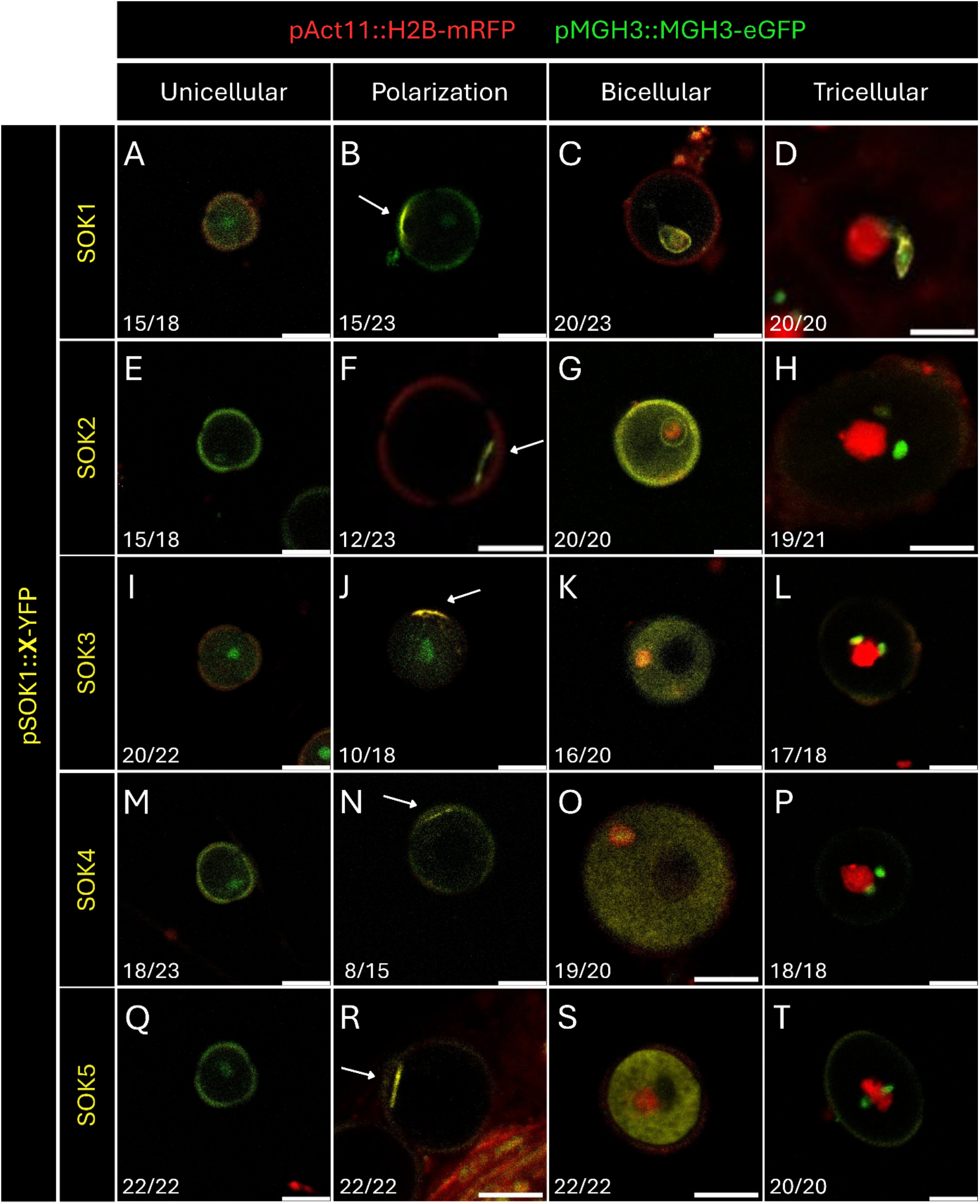
SOK2–SOK5 form an early polar domain when expressed from the SOK1 promoter but are not retained around the germline. (A–T) Confocal images of male gametophytes at the unicellular, polarization, bicellular, and tricellular stages expressing pSOK1::X-sYFP, where X is SOK1 (A–D), SOK2 (E–H), SOK3 (I– L), SOK4 (M–P), or SOK5 (Q–T). All constructs were imaged in the reporter line FB037, which expresses pACT11::H2B-mRFP (red; nuclei) and pMGH3::MGH3-eGFP (green; germline nuclei). Arrows mark polar SOKs-YFP domains. Numbers indicate pollen with the displayed pattern out of pollen scored from 4 independent lines. Scale bars, 10 µm.

### SOK1 polarity precedes, and is independent of asymmetric microspore division

SOK1 polarizes in microspores prior to nuclear migration (Fig. 2A, 3B), which has so far been the earliest sign of polarization. The nuclear migration forecasts the asymmetric PMI division (Mizuta et al., 2026; Park et al., 1998). Therefore, SOK1 polarization occurs prior to asymmetric pollen mitosis. We asked if indeed, SOK1 polarization acts upstream and independently of this event. Very few mutants are known to disturb PMI, but mutations in the microtubule-associated GEM1/MOR1 protein disturb asymmetric PMI (Park and Twell, 2001; Twell et al., 2002). GEM1/MOR1 is required for phragmoplast function (Twell et al., 2002): *gem1* microspores show incomplete nuclear migration and defective division asymmetry and cytokinesis, producing symmetric or partial divisions (Park and Twell, 2001; Park et al., 1998). We introduced SOK1-YFP into this mutant. In pollen from *gem1*/+ plants, SOK1-YFP still accumulated in a polarized domain at the cell periphery in microspores in which division had failed (Fig. 4A–B″). All transgenic lines showed at least 80% of the microspores observed displayed SOK1-YFP polar localization before PMI (Fig. 4C). SOK1 polarization can therefore be established in microspores that do not complete the normal asymmetric division. This places SOK1 polarization upstream of asymmetric division in microspores.

**Figure 4.**
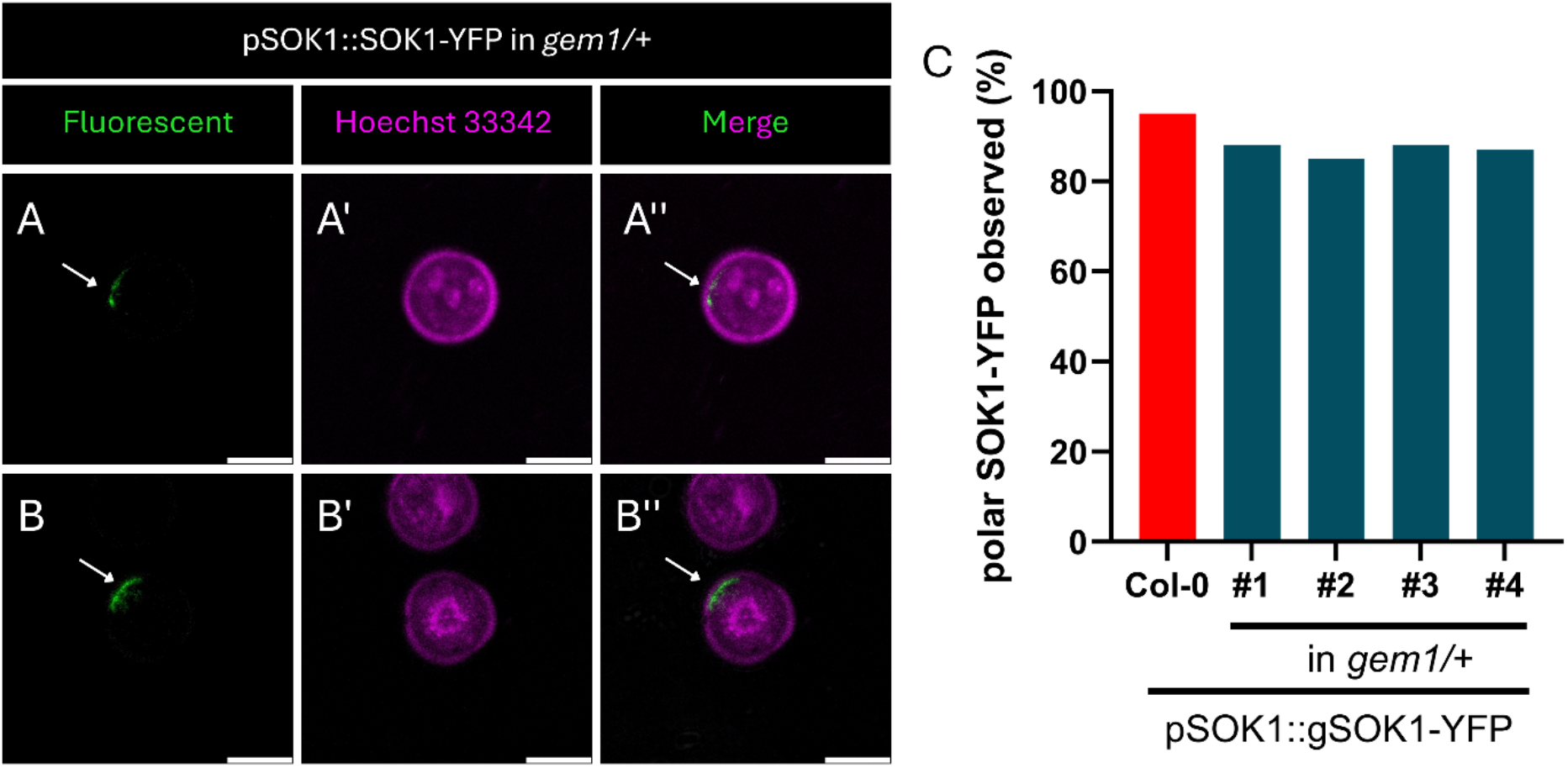
SOK1 polarization does not require completion of the asymmetric microspore division. (A–B″) Confocal sections of pollen from *gem1/+* plants expressing pSOK1::gSOK1-YFP, showing YFP fluorescence (A, B), Hoechst 33342 (A′, B′) and the merge (A″, B″). Arrows mark the polar domain. Microspores with single nucleus before pollen mitosis I were scored. (C) Percentage of unicellular microspore with polar SOK1-YFP domain, for pSOK1::gSOK1-YFP in Col-0 and four independent lines in *gem1/+* background; n = 48-50 pollens.

### SOSEKI-associated proteins collectively polarize during reproductive development

Recent, systematic proteomic analysis identified a polar protein network centered on SOSEKI proteins (Pukhovaya et al., 2026). This network includes SOSEKI proteins, as well as SOSEKI-INTERACTING BTB (SIB) proteins (SIB1/NRL2, SIB2/NRL16 and SIB3/NRL29), the cytosolic kinase SICK, the receptor-like kinases SITK1 and SITK2, and the polarized S-acyltransferases PAT19, PAT20 and PAT22 (Pukhovaya et al., 2026). An independent study placed NPT-clade NRL proteins, PATs and SICK in a lateral protein cluster in Arabidopsis roots (Yoshinari et al., 2026). Thus, SOK proteins are markers for the existence of a more general polarity system, and we explored whether other proteins in this network could inform about properties of the gametophyte cell polarity system. To this end, we analyzed the expression and localization of SIB1, SIB2, SIB3, SIB4, SIB5, SIB6, SICK, SITK1 and PAT19 in both female and male gametophytes.

In the female gametophyte at Stage I, we found a consistent pattern of inward localization for SIB1, SIB2, SICK, SITK1, and PAT19 (Fig. 5A, F, P, U, Z), similar to what was observed for SOK2, 3, and 5 (Fig. 1F, K, U). SIB3, 4, 5 and 6 are not expressed during this stage (Fig. S3). During Stage II and III SIB1, SIB2, SITK and PAT19 all show strongly overlapping expression patterns in the sporophytic tissues, generally polarized inwards to the developing gametophyte. SICK displays a mostly cytosolic localization, but is also partially polar in some cells (Fig. 5).

**Figure 5.**
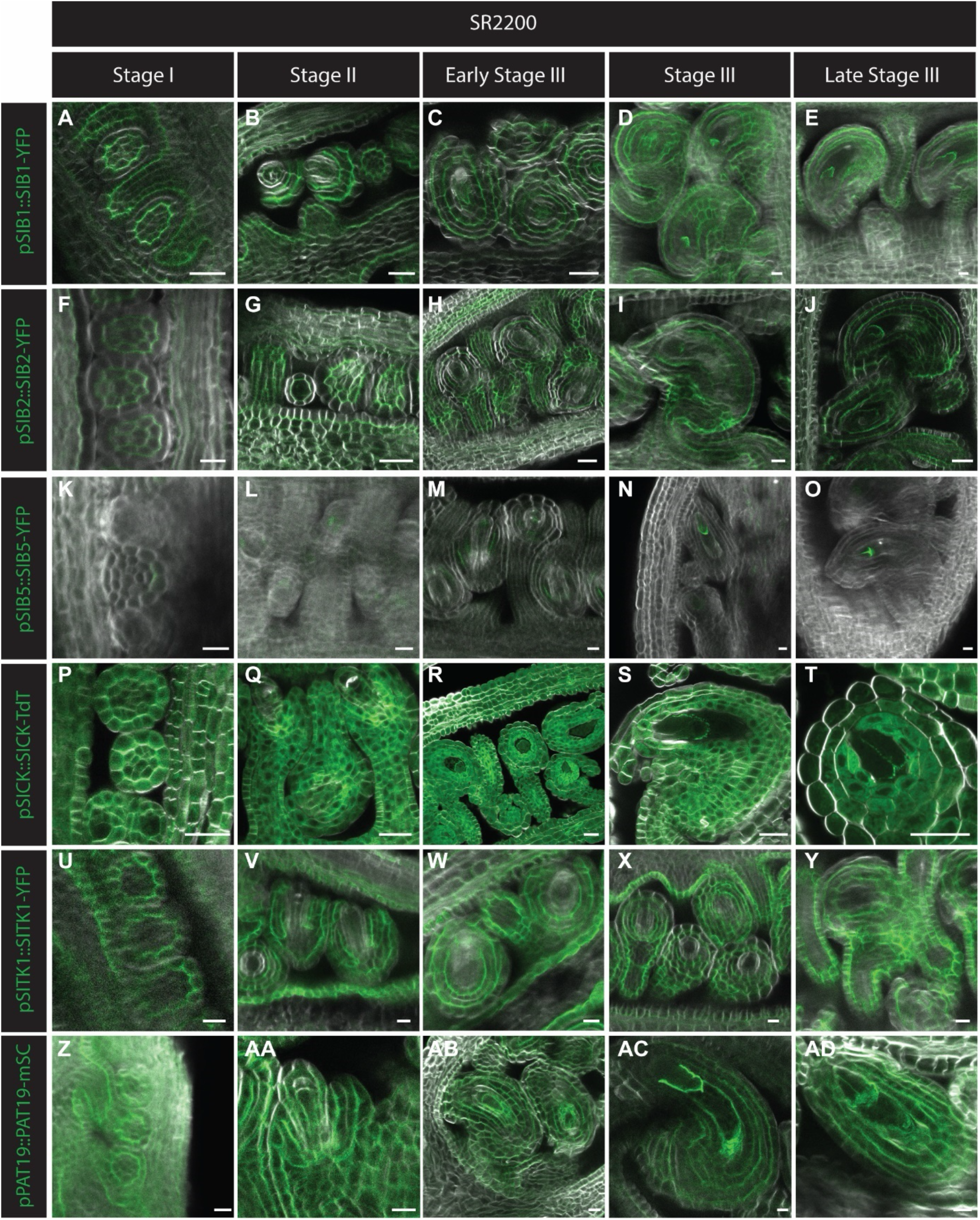
Many SOK interacting proteins are expressed and polarly localized during ovule development, only SIB5 is restricted to the female gametophyte. (A-AD) Confocal sections of the developing ovule in stage I, stage II and III expressing pSIB1::gSIB1-YFP (A-E), pSIB2::gSIB2-YFP (F-J), pSIB5::gSIB5-YFP (K-O), pSICK::gSICK-TdT (P-T), pSITK::gSITK-YFP (U-Y) and pPAT19::gPAT19-mSC (Z-AD). YFP, TdT, mSC, green; SR-2200, white. Images are displayed with brightness normalization and in some cases rotated or inverted to facilitate comparison of images. Tissues were fixed, cleared using Clearsee alpha (Kurihara et al., 2021) and stained with SR-2200 dye (Musielak et al., 2015) before imaging. Scale bars, 10 µm.

During Stage II and early Stage III, SIB1, SITK and PAT19 are polarly expressed in the forming inner and outer integuments (Fig. 5B, V, AA). During the late Stage III, SIB1, 2 and PAT19 strongly overlap within the female gametophyte, being expressed both at the chalazal end like SOK1, and in the region where the egg cell and synergids will be formed (Fig. 1B-D, 5C-E, H-J, AB-AD). SICK shows a similar pattern as SOK5 in the synergid cells (Fig. 1Y, 5T). Lastly, SIB5 is weakly expressed from Stage II onwards in the MMC/megaspore and more strongly expressed towards the end of Stage III, where it has a similar pattern as SIB1, 2, and PAT19 within the female gametophyte (Fig. 5C-E, H-J, M-O, AB-AD). Thus, SOSEKI-associated proteins show diverse patterns of polaraization that each match one or more SOSEKI proteins.

### SOSEKI-associated proteins reveal hierarchy of polarization in the male gametophyte

We next analyzed the localization of all SOSEKI-associated proteins in male gametophytes. Among the six SIB reporters, SIB1 and SIB4 reproduced the SOK1 pattern (Fig. 6A–C, J–L). Both accumulated asymmetrically at the periphery of unicellular microspores, outlined the generative cell after PMI, and surrounded the two sperm cells after PMII. SIB3, SIB5 and SIB6 could not be detected (Fig. 6G-I, M-O, P-R). SIB2 was detectable at all three stages, with cortical signal in bicellular and tricellular pollen. SICK behaved like SOK1, SIB1 and SIB4. It accumulated asymmetrically at the periphery of unicellular microspores, then surrounded the generative cell after PMI and the two sperm cells after PMII (Fig. 6S–U). PAT19 formed peripheral patches in unicellular microspores and was subsequently detected around the generative and sperm cells, although the signal was weaker in tricellular pollen (Fig. 6Y–AA). By contrast, SITK1 gave no detectable signal at any stage examined (Fig. 6V–X), in contrast to its polar localization in roots (Pukhovaya et al., 2026). SOK1, SIB1, SIB4, SICK and PAT19 therefore share a developmental sequence from an early polar cortical domain to a domain surrounding the generative and then the sperm cells.

**Figure 6.**
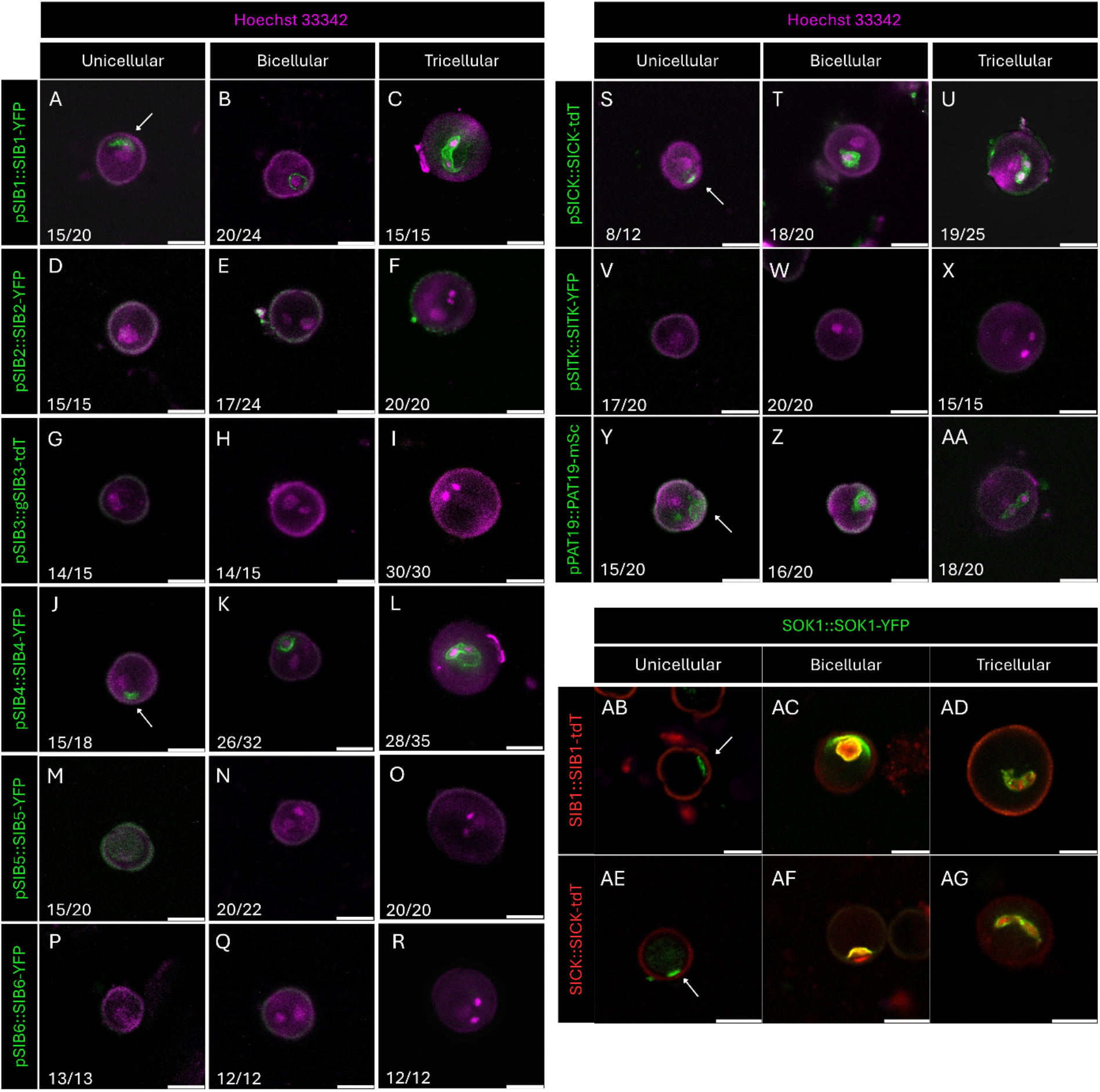
A subset of SOK-associated proteins shares the SOK1 distribution during pollen development. (A– AA) Confocal sections of unicellular microspores, bicellular pollen and tricellular pollen expressing pSIB1::gSIB1-YFP (A–C), pSIB2::gSIB2-YFP (D–F), pSIB3::gSIB3-tdT (G–I), pSIB4::gSIB4-YFP (J–L), pSIB5::gSIB5-YFP (M–O), pSIB6::gSIB6-YFP (P–R), pSICK::gSICK-tdT(S–U), pSITK1::gSITK1-YFP (V–X) and pPAT19::gPAT19-mScarlet (Y–AA). The tagged protein is shown in green in these panel — note that SICK-tdTomato and PAT19-mScarlet are red-emitting fluorophores displayed in green for consistency with the YFP reporters. (AB–AG) Confocal sections of pollen co-expressing pSOK1::gSOK1-YFP with pSIB1::gSIB1-tdTomato (AB–AD) or pSICK::gSICK-tdTomato (AE–AG), at the unicellular, bicellular and tricellular stages. SOK1-YFP, green; SIB1-tdTomato and SICK-tdTomato, red. Hoechst 33342, magenta. Arrows mark polar cortical domains. Numbers indicate pollen with the displayed pattern, scored from four independent lines. Scale bars, 10 µm

Having multiple proteins with similar polar patterns in the same, simple structure, we next asked if there is an order in which proteins polarize. We therefore crossed the SOK1-YFP line with SICK-tdTomato or SIB1-tdTomato lines and co-localized the two fluorescent proteins. In both cases, we found that SOK1 protein polarized before either of the other two proteins (Fig. 6AB, AE). As suggested by individual patterns (Figs. 2A-B, S-T), each of the two pairs ultimately co-localized to the microspore polar crescent (Fig. 6AC, AF) and the PGCM (Fig. 6AD, AG). This suggests that SOK1 likely acts prior to SIB1 and SICK in the polarity network.

### A universal polarity system guides localization in Arabidopsis female gametophytes

The analysis of 14 polar proteins (5 SOK proteins, 6 SIB proteins, SICK, SITK1 and PAT19) reveals novel polar cellular domains in both male and female gametophytes. However, each protein marks a subset of these polar domains. Misexpression of all SOK proteins in the SOK1 domain shows that other proteins either localize to the SOK1 domain, or remain cytosolic. This suggests that perhaps there are no alternative polarity domains at this stage. To test this directly, we expressed SOK1 ubiquitously from the *RPS5A* promotor (Weijers et al., 2001) and analyzed localization in both male and female gametophytes. In male gametophytes, this line mirrored native SOK1 accumulation (Fig. 2A-C, 7A-C). In female gametophytes, SOK1 strongly overlapped with SIB1, 2, SITK and PAT19 (Fig. 5C-E, H-J, M-O, AB-AD, 7D-G). Strikingly, SOK clearly localized the inner membrane of synergid cells (Fig 7G). This is very similar to native SOK5 localization, as well as patterns of SICK, SIB5, and PAT19 (Fig. 1Y, 5O, T, AD). This data suggests that these proteins collectively report on a single polarity system.

**Figure 7.**
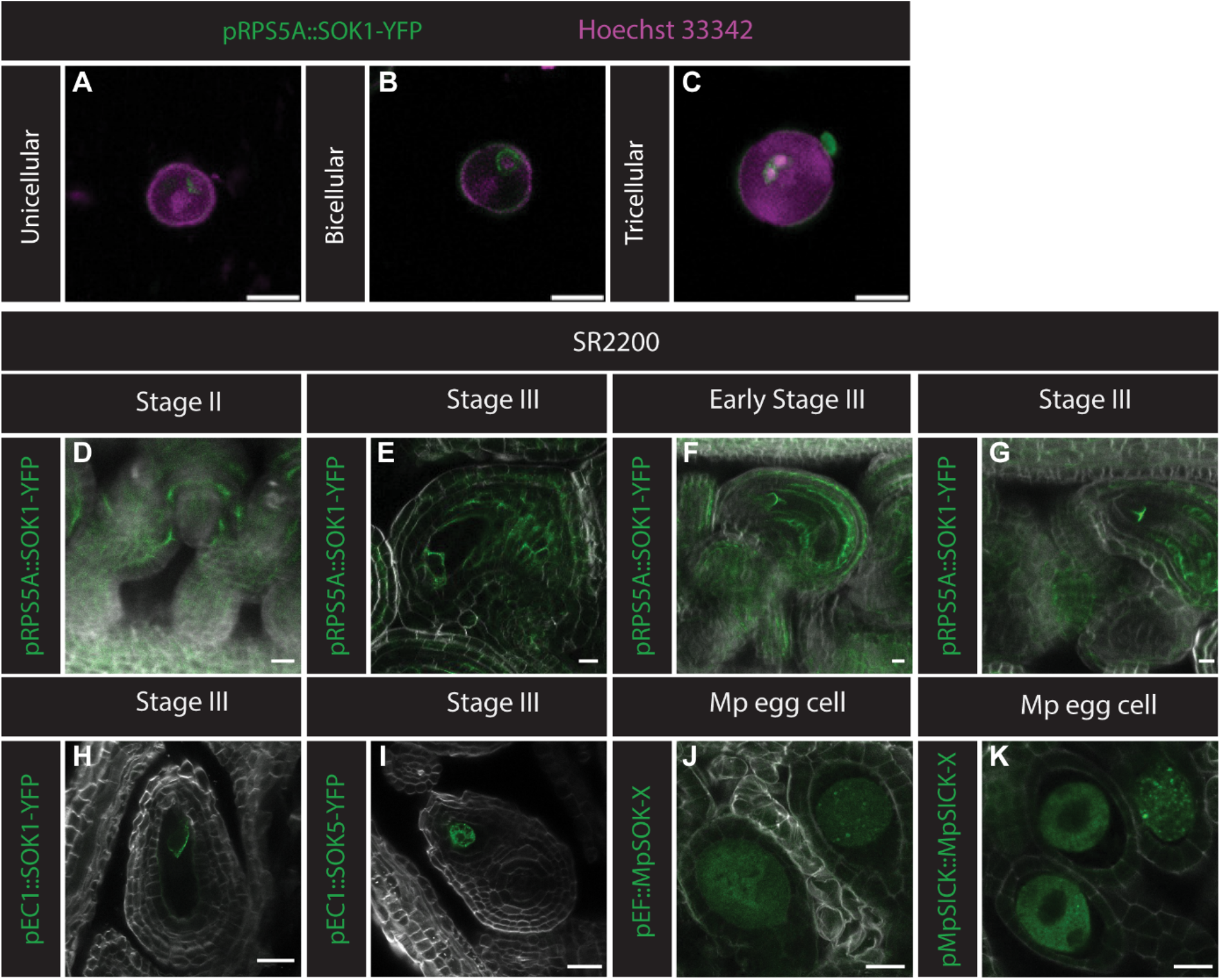
Misexpression of SOK1 shows that the described polar complex is one system not multiple, furthermore egg cell polarity is not conserved during evolution. (A-K) Confocal sections of the developing microspores, bicellular, and tricellular pollen and ovule in stage II and III expressing pRPS5A::gSOK1-YFP. (A-G) Confocal sections of ovules in late stage III expressing pEC1::SOK1-YFP and pEC1::SOK5-YFP (H, I). Confocal sections of Marchantia egg cells, expressing EF::MpSOK-YFP and MpSICK::MpSICK-YFP (J, K). YFP, TdT, mSC, green; SR-2200, white; Hoechst 33342, magenta. Images are displayed with brightness normalization and in some cases rotated or inverted to facilitate comparison of images. Tissues were fixed, cleared using Clearsee alpha (Kurihara et al., 2021) and stained with SR-2200 dye (Musielak et al., 2015) before imaging. Scale bars, 10 µm.

**Figure 8.**
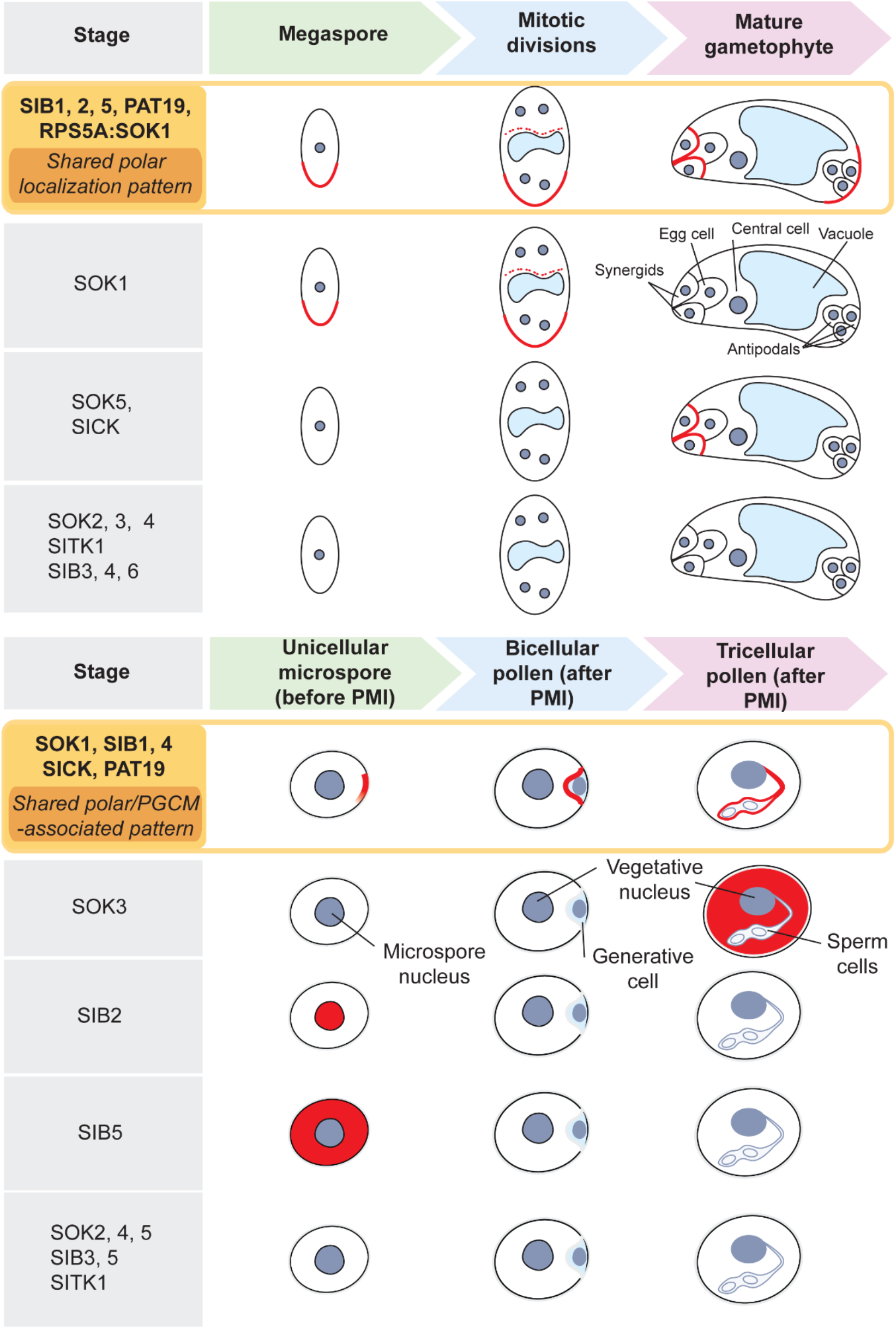
Visual summary of the main findings of the paper.

We next asked if cells in which we did not find native expression of SOSEKI proteins can polarize the proteins. We therefore expressed SOK1 and SOK5 from the egg cell-specific *EC1* promotor (Sprunck et al., 2012). In these lines, SOK1 polarized towards the antipodal end of the egg cell in a similar direction as under the native promotor, confirming that the pathway that polarizes SOK1 is present in the egg cell (Fig. 7H). Interestingly, SOK5 is mostly cytosolic and does not show any polarized expression, even though it is polarly expressed in the synergids when expressed from its native promotor (Fig. 7I). Therefore, this data suggests that while there is a common polarity system, SOK1 and SOK5 differ in their capacity to be polarly targeted, similar to the case in pollen development.

We previously found that endogenous SOK proteins are polarized in gametophyte tissue in the liverwort Marchantia and the moss Physcomitrium (Pukhovaya et al., 2026; van Dop et al., 2020). We however did not explore polarization in gametes. The finding that the Arabidopsis egg cell is capable of polarizing SOK1 urges the question of whether, likewise, bryophyte egg cells polarize proteins in the SOK network. We studied the localization of Marchantia SOK (MpSOK) and SICK (MpSICK) proteins in Marchantia egg cells. Unlike Arabidopsis, Marchantia produces large, spherical egg cells that are only covered by a single layer of cells. We expressed MpSOK and MpSICK from the ubiquitous *EF* promotor and the endogenous promotor, respectively. In these lines, MpSOK is completely cytosolic in egg cells and shows a punctate pattern (Fig. 7J). Likewise, MpSICK is also cytosolic, and also shows punctate pattern (Fig. 7K). This data suggest that, in contrast to the Arabidopsis egg cell, the Marchantia egg cell does not polarize SOK and associated proteins, and may not be not polarized.

## Discussion

Land plants have remarkable life cycles that alternate multicellular haploid and diploid generations (Hofmeister, 1851). From an ancestral early land plant, two lineages emerged with nearly opposing life histories regarding the role of each of the generations: Bryophytes feature a dominant, and indeterminately growing haploid (gametophyte) phase and a generally diminutive diploid (sporophyte) phase that is transient and serves to form spores following meiosis (Shaw et al., 2011). With the exception of ferns, Tracheophytes have a dominant and indeterminate sporophyte and a diminutive gametophytes (Kenrick, 2000) that divides a few times to generate gametes (Maheshwari, 1950). Bryophytes gametophytes and Tracheophyte sporophytes share complex multicellular development, patterning and meristem activity, but these are entirely different phases of life and an important question is whether the same mechanisms of development underlie both, how these relate to the other generation within each lineage, and how these derive from the ancestral land plant. Here we explored the homology of cell polarity systems in the generations within the flowering plant Arabidopsis. There has been a rich documentation of cell polarity in the sporophyte (Muroyama and Bergmann, 2019), but so far, no markers for cell polarity have been reported in the development of their gametophytes. Our work was inspired by the recent discovery of deeply conserved cell polarity proteins, including SOSEKI (van Dop et al., 2020; Yoshida et al., 2019), SOSEKI-INTERACTING BTB (SIB), SIB1-INTERACTING CYTOSOLIC KINASE (SICK) and PROTEIN ACYLTRANSFERASE19 (PAT19) (Pukhovaya et al., 2026; Yoshinari et al., 2026), that are polarized both in Arabidopsis sporophytes and Marchantia gametophytes. This suggests that a common polarity system me be shared by the two generations across these species, and by inference, perhaps also within each species. Given that both male and female gametophytes in Arabidopsis undergo multiple divisions, including asymmetric ones (Borg and Twell, 2010), and the female gametophyte grows anisotropically (Drews and Yadegari, 2002), it is likely that cell polarity plays a role in gametophyte development.

We report that both male and female gametophytes are marked by early polar localization of several proteins. In fact, the polarization of SOK1 marks symmetry breaking in microspore well before any other sign of asymmetry emerges. In the female gametophyte SOK1 likewise decorates one pole of the surviving megaspore prior to expansion growth. Hence, establishment of cell polarity is indeed an early event in the Arabidopsis haploid phase. Following this initial polarization, there are addition polarization events in both gametophytes: in the male, there is polar targeting of a range of proteins to the PGCM, the membrane that surrounds the generative cell and sperm (Sugi et al., 2024). In the female, there is polarization within the two synergid cells, and there is the capacity of the egg cell to polarly localize ectopically expressed SOSEKI protein. Hence, in both gametophytes there are at least two polar domains, and perhaps multiple steps in polarization. In the case of the male, we could dissect the independence of the initial polarization to a polar crescent in the microspore and the later targeting to the PGCM. Ectopically expressing all SOK proteins from the SOK1 promoter revealed that all SOK proteins can mark the polar crescent, while only SOK1has the ability to localize to the PGCM. This is interesting, because the PGCM is contiguous with the initial polar crescent. Since the crescent marks the region where the asymmetric division forms, the crescent essentially “engulfs” the generative cell. Yet, the properties of this membrane must change during the process, given the loss of SOK2/3/4/5 from it. Detailed analysis of SOK1 localization during the first asymmetric microspore division through live imaging should help resolve whether the polar domain is indeed inherited to form the PGCM, or whether there is dynamic rearrangement of membrane and proteins during the process.

In the female, while there are multiple polar domains, these occur in different cells (megaspore versus synergids versus egg cell), and there is no evidence that within the same cell, there are different capacities to polarize, as in the male. Indeed, when ectopically expressing SOK1, it was able to mark all polar domains discovered through the native expression of the other proteins studied. We interpret that there is a single polarity system in the female gametophyte that guides the localization of all the proteins we tested.

It is remarkable that the full set of conserved polarity-localized proteins (SOK, SIB, SICK, PAT) localize to the same polar domains in gametophytes. This suggests that this set of proteins may indeed represent a module that is functionally interlinked. So far, it has been difficult to define the interdependencies within this module, given that there is such strong co-localization in the growing sporophyte. The gametophytes offer an exquisite context for studying causalities, given that there is a linear, directional ontogeny where one stage precedes the next. We co-localized SOK1 and SIB1, as well as SOK1 and SICK, and this revealed that SOK1 appears to accumulate at the polar crescent in microspores before the other two proteins. While we cannot exclude that there are functionally relevant polarized pools of SIB1 and SICK contemporaneously or preceding SOK1 that are below the detection limit, the simplest interpretation is that SOK1 acts upstream of SIB1 and SICK in microspore polarization. Detailed analysis of mutants in each of these components should resolve such functional relationships. In addition, there are enzyme-substrate relationships between some components, such as palmitoylation of SOK1 by PAT19 (Pukhovaya et al., 2026), that lead to testable hypotheses, such as that a Cysteine mutation in SOK1 should interfere with those processes that require PAT19. Analysis of mutants is also critical to resolve the function of these polarity proteins, and cell polarity in general, in the haploid generation. It is intuitive that asymmetric cell division, anisotropic growth and membrane differentiation (e.g. PGCM) would depend on polar targeting, but so far, our work only reports on the existence of polarity, not its relevance. It is encouraging that the *pat19;20;22* triple mutant has strong fertility defects (Liu et al., 2024) that may be connected to gametophyte polarity defects. Unfortunately, genetic analysis of the SOK family has do far not led to obvious phenotypes (Yoshida et al., 2019), but this may be confounded by redundancies, and perhaps even the very gametophytic localizations we report here. It is possible that there are complex requirements for SOK proteins in the haploid phase, which would complicate the generation of mutants.

A major inspiration for exploring Arabidopsis gametophyte polarity was the identification of conserved polar proteins between the Marchantia gametophyte and the Arabidopsis sporophyte. Strikingly though, we found that neither MpSOK nor MpSICK are polarized in the Marchantia egg cell. This is in contrast to AtSOK1, which does polarize in the Arabidopsis egg cell when expressed there. There are several potential explanations: first, it is possible that MpSOK requires geometric cell edges, which is where nearly all SOK localization is found. The capacity to break symmetry in a spherical cell may be an evolutionary innovation in AtSOK1. This is perhaps unlikely, given that all AtSOK proteins can break symmetry when expressed in microspores. Alternatively, the Marchantia egg cell may simply not polarize. The essentially symmetric division of the zygote (Kohchi et al., 2021) is consistent with this notion. It is clear that the availability of molecular markers of cell polarity now allows asking and answering new questions related to the evolution of polarity and the way it is integrated in multicellular development.

More generally, while the current study describes the existence of a cellular and molecular polarity system in flowering plant gametophytes, we consider this the starting point for a range of previously inaccessible roads. First, using these novel markers for more detailed and dynamic descriptions of cell polarity will lead to a solid, quantitively and hopefully harmonizing framework for sporophyte and gametophyte development across the Bryophyte-Tracheophyte divide. Second, this work postulates roles for polarity proteins in gametophyte development, which can now be tested. Third, these proteins – as markers – will help to place cell polarization in the known frameworks of gametophyte development, as we did with the *gem1* mutant. Finally, it should be clear that the rich polarity landscape in both gametophytes, and their potential connection to formative developmental events, suggests that gametophyte polarization may be a developmental bottleneck, and may explain why mutagenesis and screening at post-embryonic stages have so far produced so few general cell polarity regulators in plants. We propose that gametophyte cell polarization offers a potentially fertile ground for genetic exploration of plant cell polarity mechanisms in the future.

## Materials and methods

### Plant material and growth conditions

The *Arabidopsis thaliana* Columbia-0 (Col-0) accession was the wild-type background for all reporter lines produced in this study. The vegetative and gamete lineage reporter system FB037 containing both pHTR10:HTR10:eGFP and pACT11:H2B:mRFP has been described previously (Borges et al., 2012). Seeds were surface-sterilized and then stratified for 2 days in the dark at 4°C. Seeds were then germinated and grown under continuous white light at ∼120 μmol⋅m^−2^⋅s^−1^ and 22°C on vertically placed Petri dishes containing 0.5× Murashige and Skoog (MS) medium (Murashige and Skoog, 1962) supplemented with 1% agar and 0.3% sucrose. The seedlings were then transferred to soil and grown at 22 °C in long-day conditions (16 h light / 8 h dark) in controlled-environment rooms, unless otherwise specified. Phenotypic analysis was always carried out with pollen from at least four different plants.

### Vector construction and generation of transgenic lines

The p*SOK1::SOK-YFP, pSOK2::SOK2-YFP, pSOK3::SOK3-YFP, pSOK4::SOK4-YFP, pSOK5::SOK5-YFP and pRPS5A-SOK1-YFP* transgenic lines have previously been described (Yoshida et al., 2019). The *pSIB1::SIB1-YFP*, *pSIB2::SIB2-YFP*, *pSIB3::SIB3-TdTomato, pSICK::SICK-TdTomato* and pSITK1::SITK1-YFP have been described (Pukhovaya et al., 2026). Constructs for YFP-tagged SOK1-paralogs driven by SOK1 promoter were created using the Golden Gate modular cloning method (Engler et al., 2014). Briefly, The SOK1 promoter (2004bp upstream of the start codon of SOK1) was amplified and cloned into pICH41295. The genomic coding sequence of all SOK paralogs were amplified and cloned into pICSL01005. To generate the level 1 module, the SOK1 promoter, individual SOK paralog, a YFP-tag (cloned into pAGM1301) and the NOS terminator (pICH41421) were combinatorially assembled into the Level 1 acceptor backbone (pICH47742). For the final Level 2 assembly, each SOK Level 1 module was assembled into the Level 2 acceptor backbone (pICSL4723) with a FAST-Red selectable marker (pICSL11015) and linker Ele2 (pICH41744). To generate SIB4, SIB5 and SIB6 reporter lines, sequences containing the relevant promoter and coding sequence were amplified and assembled upstream of sYFP and NOS terminator sequences in HpaI-digested pPLV17 (De Rybel et al., 2011) using HiFi assembly (primers contain 20 bp tails to facilitate homology-mediated assembly to neighboring fragments). A GGGSGGGGS flexible linker was introduced between each coding sequence and sYFP. To generate pEC1::SOK1-YFP and pEC1::SOK5-YFP, coding sequence SOK1 and SOK5, excluding the stop codon, were cloned into linearized pDONR221 using NEBuilder® HiFi DNA Assembly (New England Biolabs). The pDONR221 vector was amplified with the primers 5′-GAAGCCTGCTTTTTTGTACAAAGTTGGC-3′ and 5′-GACCCAGCTTTCTTGTACAAAGTTGGC-3′. The resulting entry clones were subsequently recombined into a modified pMpGWB destination vector via an LR reaction using Gateway® LR Clonase® II Enzyme Mix (Thermo Fisher Scientific). In this destination vector, a YFP cassette had been inserted into the SacI restriction site of pMpGWB101, enabling the generation of C-terminal YFP fusions of the target proteins. To drive expression in egg cells, the EC1 promoter fragment was cloned into the HindIII restriction site, replacing the original EF1α promoter present in pMpGWB101. Primers used for generation of these constructs are listed in Supplemental Table S1. All constructs were introduced into *Agrobacterium tumefaciens* strain GV3101pMP90 and transformed using the floral dip method (Clough and Bent, 1998). The MpSOK and MpSICK transgenic Marchantia lines were previously described (Pukhovaya et al., 2026).

### Confocal Imaging and Analysis

Leica TCS SP5 confocal microscope was used with the following fluorescence excitation emission setting to visualize reporter genes and staining signals: SR2200 and Hoechst 33342 excitation at 405 nm, emission 420 nm, YFP excitation 515 nm, emission 527 nm; eGFP excitation 488 nm, emission 507 nm; RFP excitation 584 nm, emission 607 nm. Images were taken with 20x water immersion objectives. For analysis, Fiji and Las X image analysis software were used (Schindelin et al., 2012). Brightness was adjusted uniformly across the entire image, and identical adjustments were applied when signal intensity was directly compared.

### Imaging the microspore

To detect fluorescent signals, anthers containing microspores and mature pollen were dissected from fresh inflorescences in 0.3 m Mannitol solution. For nuclear staining, 0.1% Triton X-100 with 0.1% Hoechst 33342 were added to the Mannitol solution.

### Imaging the ovules and Marchantia egg cells

Flowers were harvested in 4% PFA in PBS-T and fixed for 30 minutes under vacuum and incubated at 5 degrees overnight. The next day, flowers were washed 3 times in PBS-T before incubation in Clearsee-alpha (Kurihara et al., 2021) for approximately 2 weeks. During this time, Clearsee-alpha solution was refreshed regularly. Afterwards, samples were washed using PBS-T and incubated for 30 minutes under vacuum with 0.1% SR2200 (Musielak et al., 2015) in PBS-T. For imaging, samples were washed once more in PBS-T and mounted in Vectashield anti fading solution (Vector Labs). This protocol was adapted from (Vijayan et al., 2021).

## Acknowledgements

The authors thank Prof. David Twell for the *gem1* seeds, Jorg Becker for the FB037 reporter system and our team members for helpful discussions.

## Authorship statement

Conceptualization: S.K.W.H., M.v.S., C.A., D.W..; Validation: S.W.; Formal analysis: S.K.W.H., M.v.S.; Investigation: S.K.W.H., M.v.S., B.V.; Resources: C.A., J.R.R.; Writing - original draft: S.K.W.H., M.v.S.; Writing - review & editing: S.K.W.H., M.v.S., C.A., J.R.R., D.W.; Visualization: S.K.W.H., M.v.S.; Supervision: D.W.; Project administration: D.W.; Funding acquisition: S.K.W.H., M.v.S., D.W..

## Data and resource availability

All primary data is available upon request from the corresponding author.

## AI Statement

No AI was used in the preparation of this paper

