## Supplemental Figure for "The cell polarity landscape in Arabidopsis male and female gametophytes"

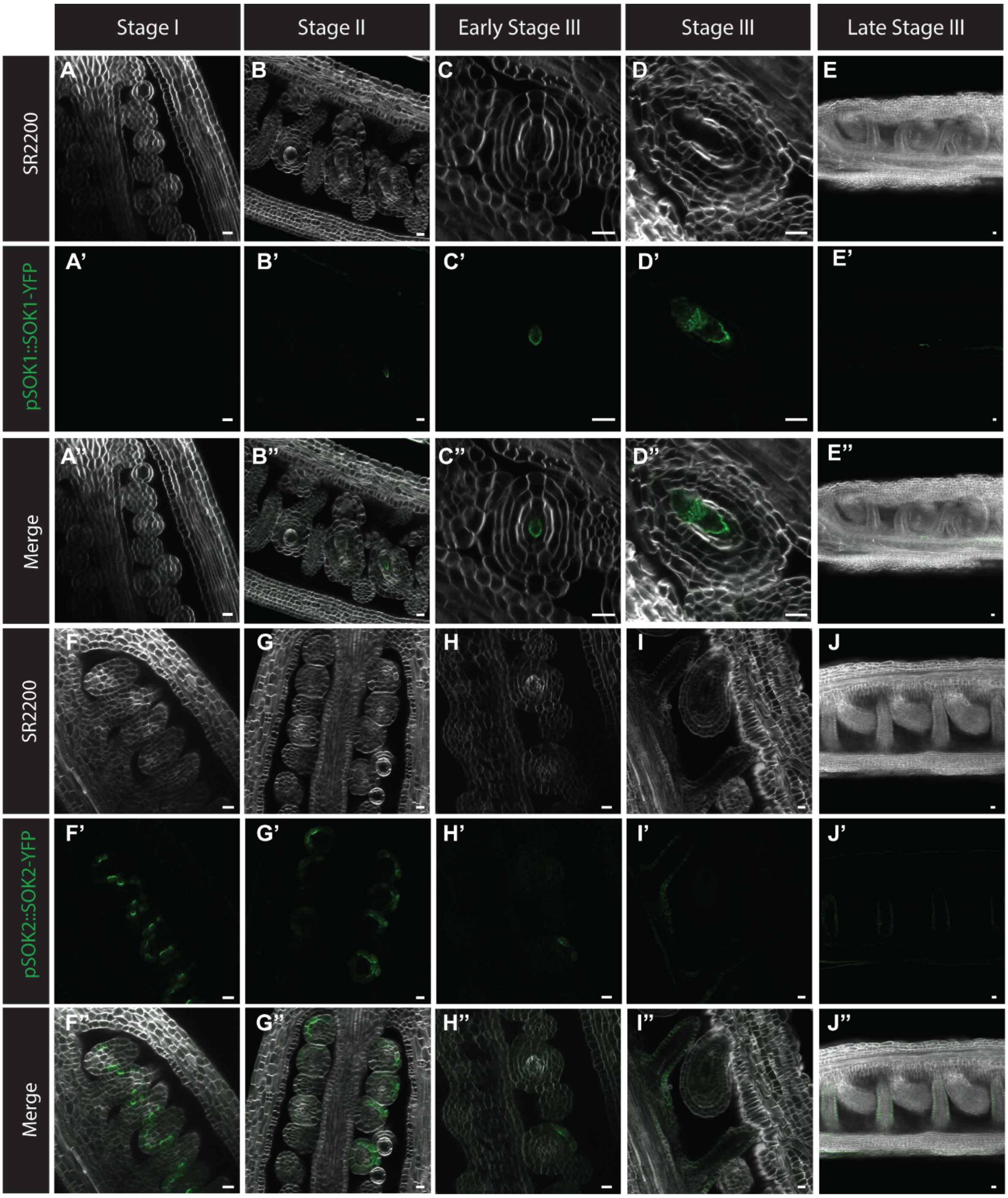

Stage I
Stage II
Early Stage III
Stage III
Late Stage III
SR2200
pSOK1::SOK1-YFP
Merge
SR2200
pSOK2::SOK2-YFP
Merge
A
B
C
D
E
A'
B'
C'
D'
E'
A''
B''
C''
D''
E''
F
G
H
I
J
F'
G'
H'
I'
J'
F''
G''
H''
I''
J''

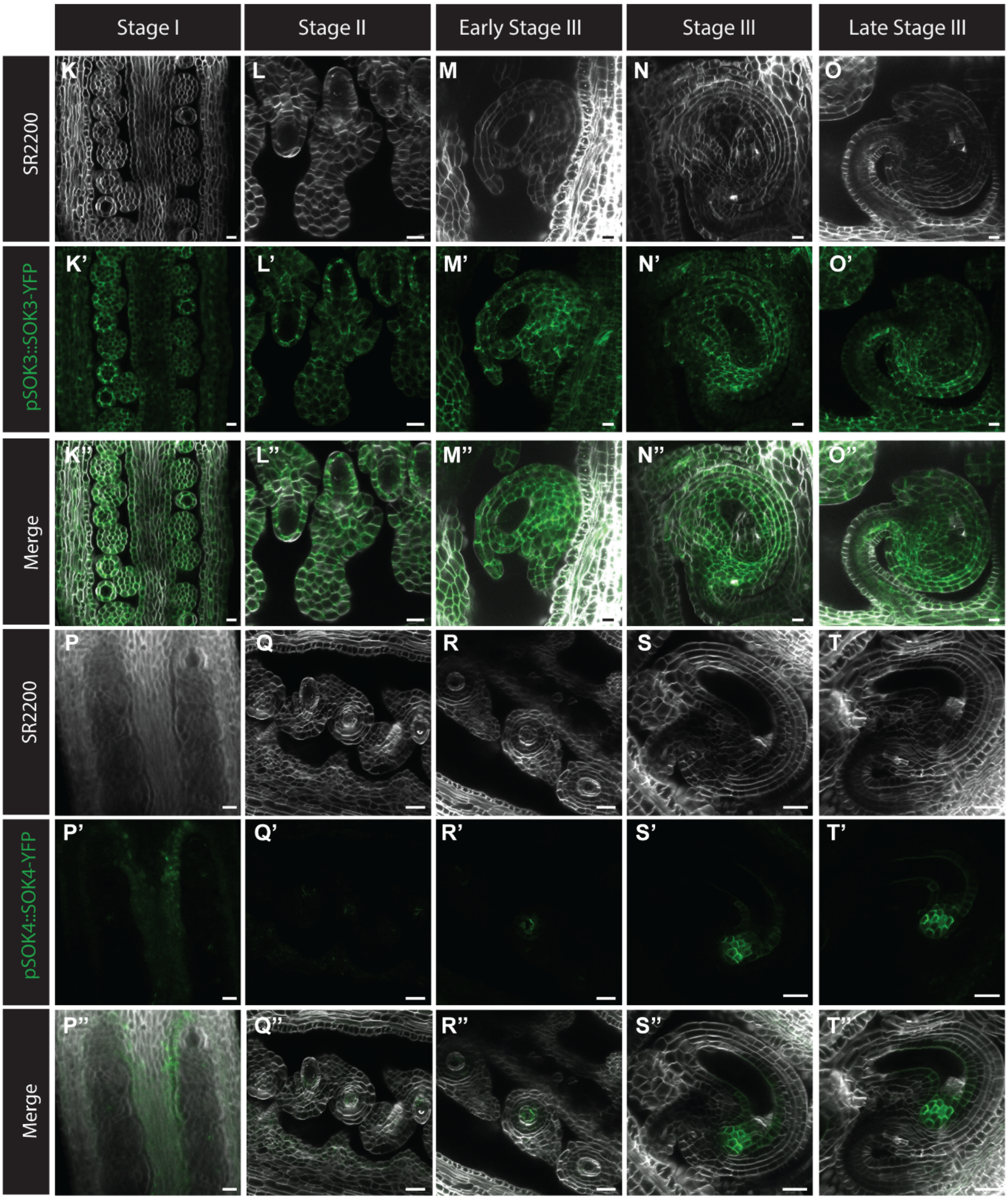
Stage I
Stage II
Early Stage III
Stage III
Late Stage III
SR2200
pSOK3::SOK3-YFP
Merge
SR2200
pSOK4::SOK4-YFP
Merge
K
L
M
N
O
K’
L’
M’
N’
O’
K’’
L’’
M’’
N’’
O’’
P
Q
R
S
T
P’
Q’
R’
S’
T’
P’’
Q’’
R’’
S’’
T’’

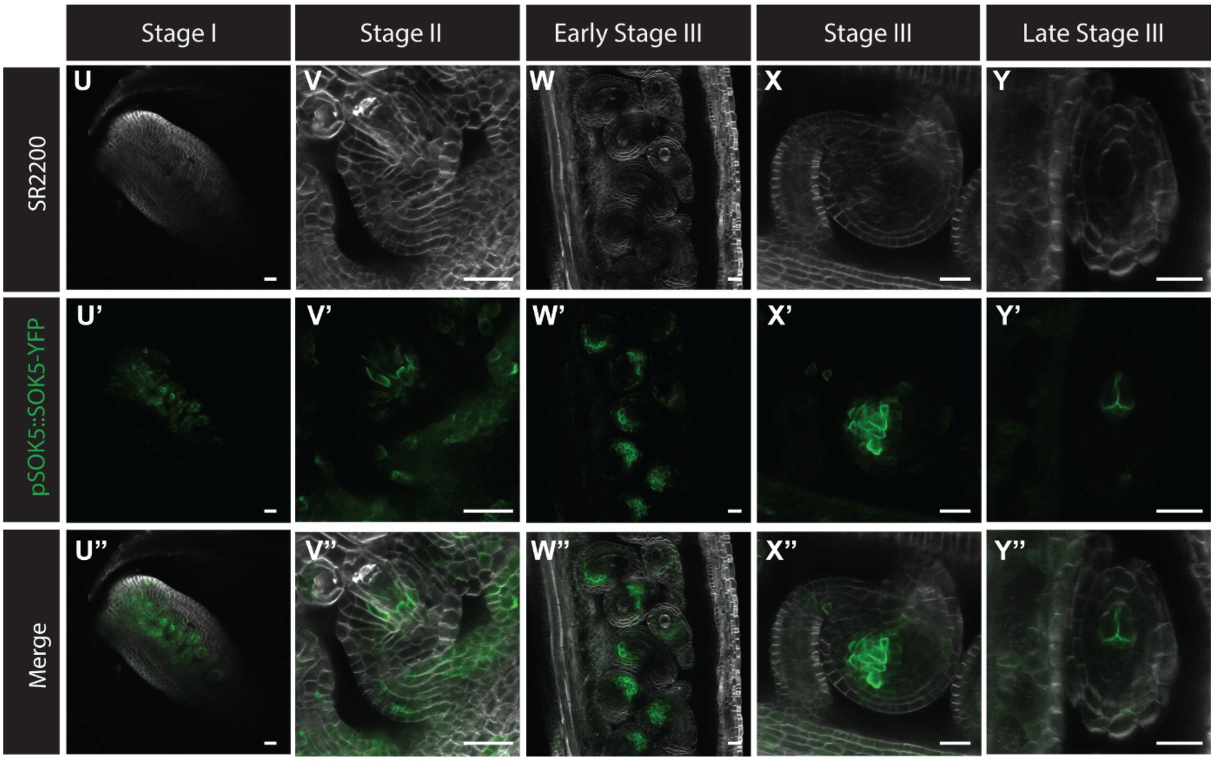


**Figure S1 Expression of SOK1-5 in the female gametophyte.** (A-Y) Confocal sections (seperated channels and merged images) of the developing ovule in stage I, stage II and III expressing pSOK1::gSOK1-YFP (A-E), pSOK2::gSOK2-YFP (F-J), pSOK3::gSOK3-YFP (K-O), pSOK4::gSOK4-YFP (P-T) and pSOK5::gSOK5-YFP (U-Y). YFP, green; SR-2200, white. Images are displayed with brightness normalization and in some cases rotated or inverted to facilitate comparison of images. Tissues were fixed, cleared using Clearsee alpha (Kurihara et al., 2021) and stained with SR-2200 dye (Musielak et al., 2015) before imaging. Scale bars, 10 µm.

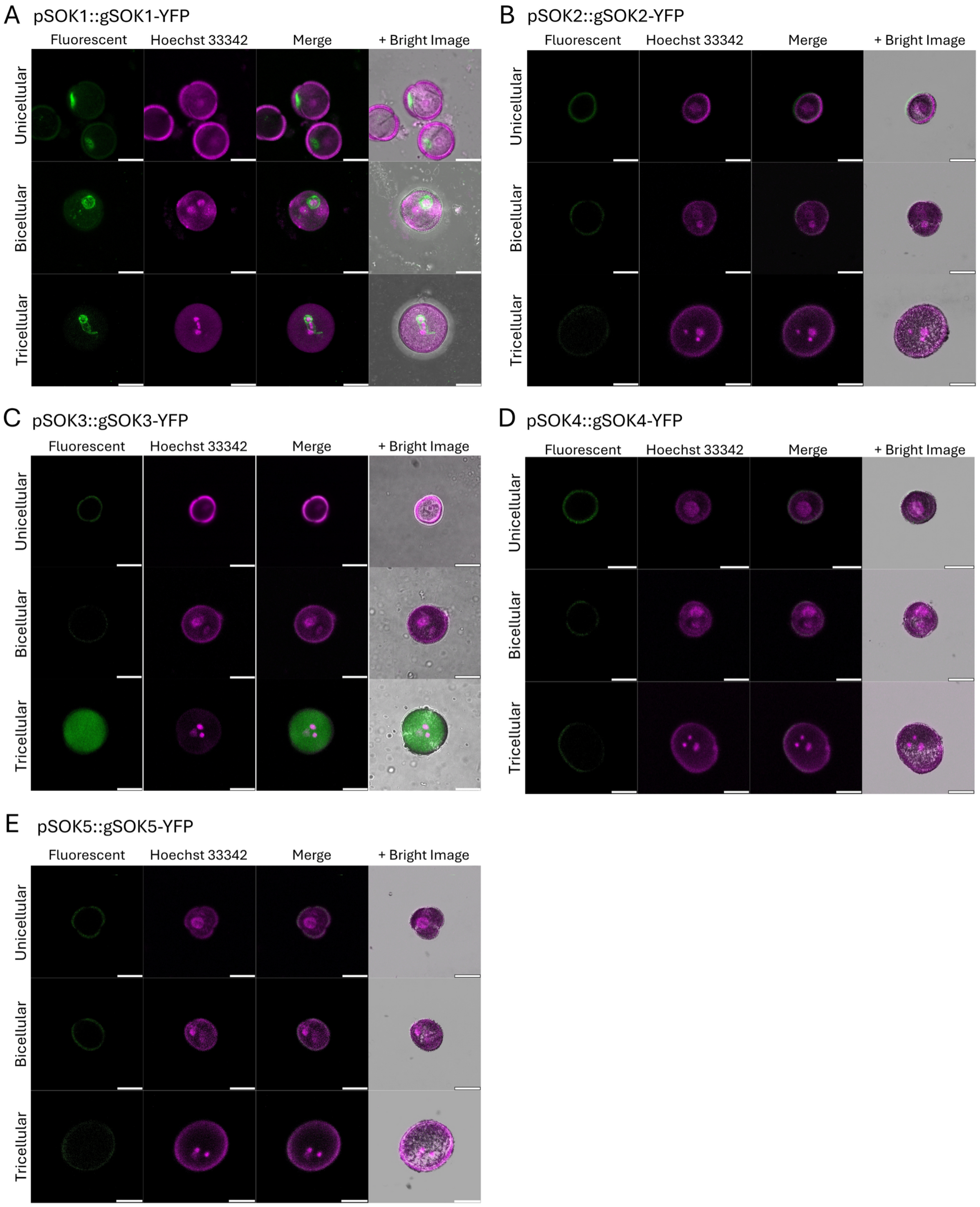


**Figure S2. Separated fluorescence channels for SOSEKI reporters in the male gametophyte.** (A–E) Confocal sections of unicellular microspores, bicellular pollen and tricellular pollen expressing pSOK1::gSOK1-YFP (A), pSOK2::gSOK2-YFP (B), pSOK3::gSOK3-YFP (C), pSOK4::gSOK4-YFP (D) and pSOK5::gSOK5-YFP (E), shown as separated channels. Within each block, rows are the three developmental stages and columns are YFP fluorescence, Hoechst 33342, the two-channel merge, and the merge overlaid on the brightfield image. YFP, green; Hoechst 33342, magenta. Scale bars, 10 µm.

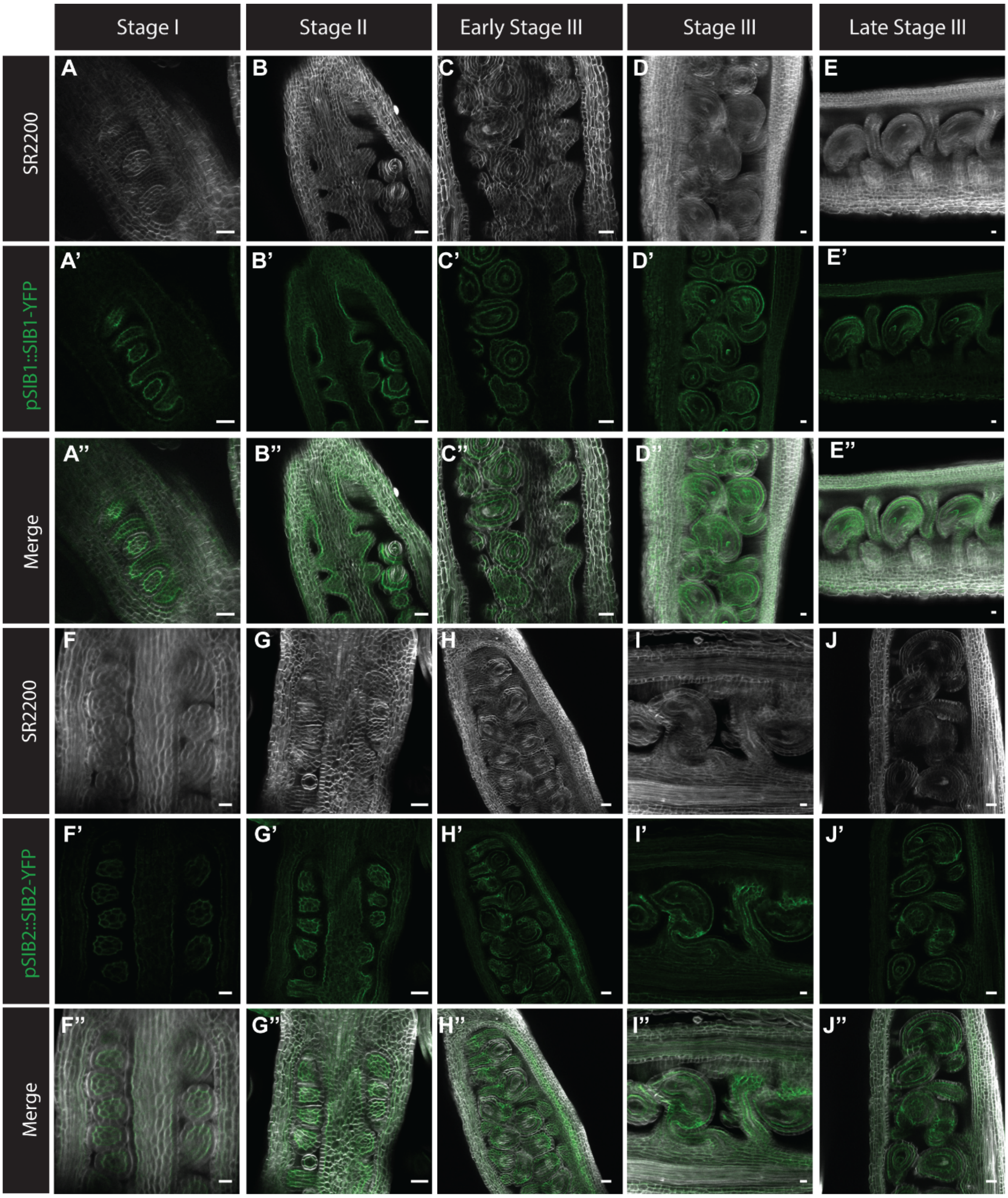

Stage I
Stage II
Early Stage III
Stage III
Late Stage III
SR2200
pSIB1::SIB1-YFP
Merge
SR2200
pSIB2::SIB2-YFP
Merge
A
B
C
D
E
A'
B'
C'
D'
E'
A''
B''
C''
D''
E''
F
G
H
I
J
F'
G'
H'
I'
J'
F''
G''
H''
I''
J''

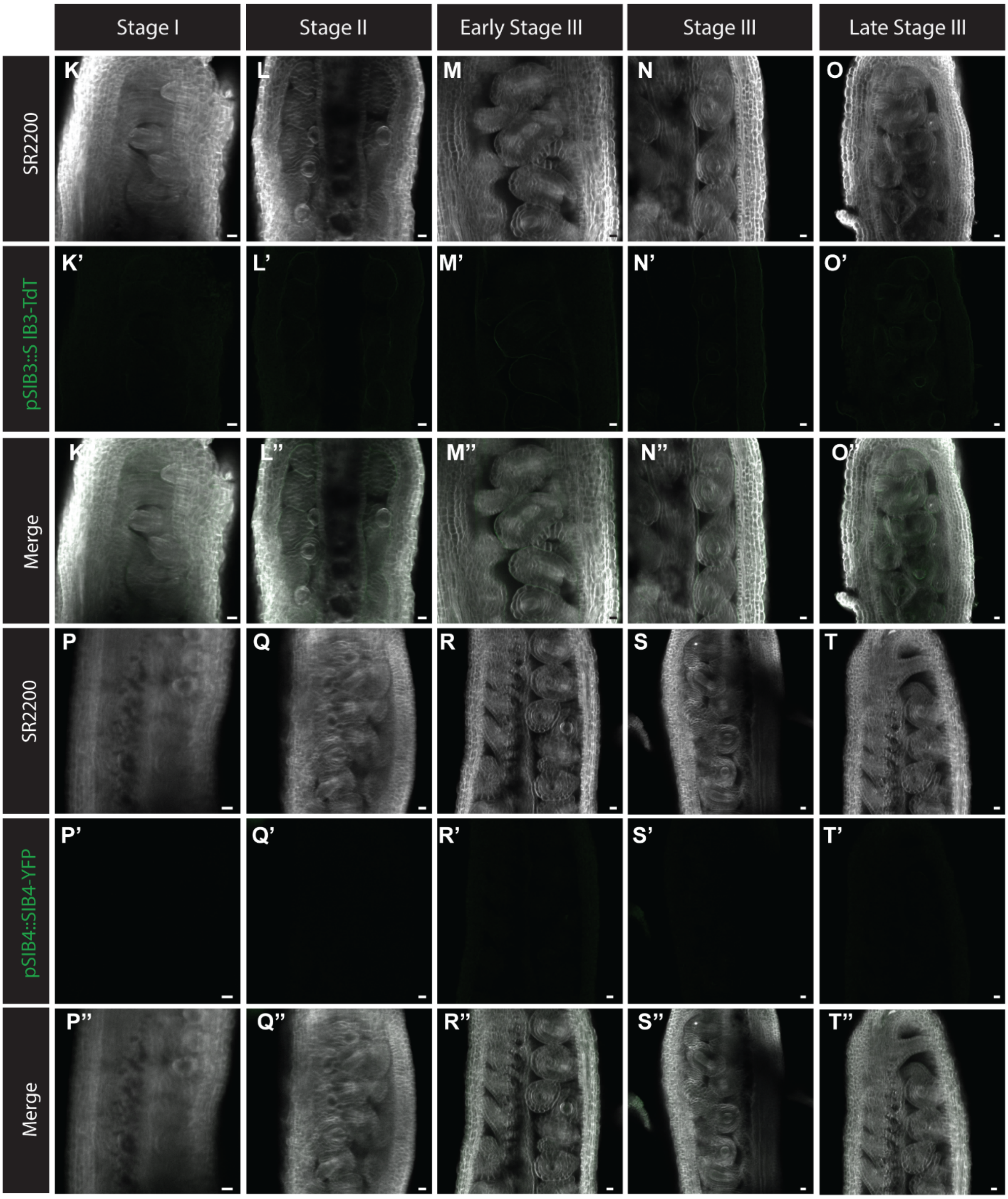
Stage I
Stage II
Early Stage III
Stage III
Late Stage III
SR2200
pSIB3::SIB3-TdT
Merge
SR2200
pSIB4::SIB4-YFP
Merge
K
L
M
N
O
K'
L'
M'
N'
O'
L''
M''
N''
P
Q
R
S
T
P'
Q'
R'
S'
T'
P''
Q''
R''
S''
T''

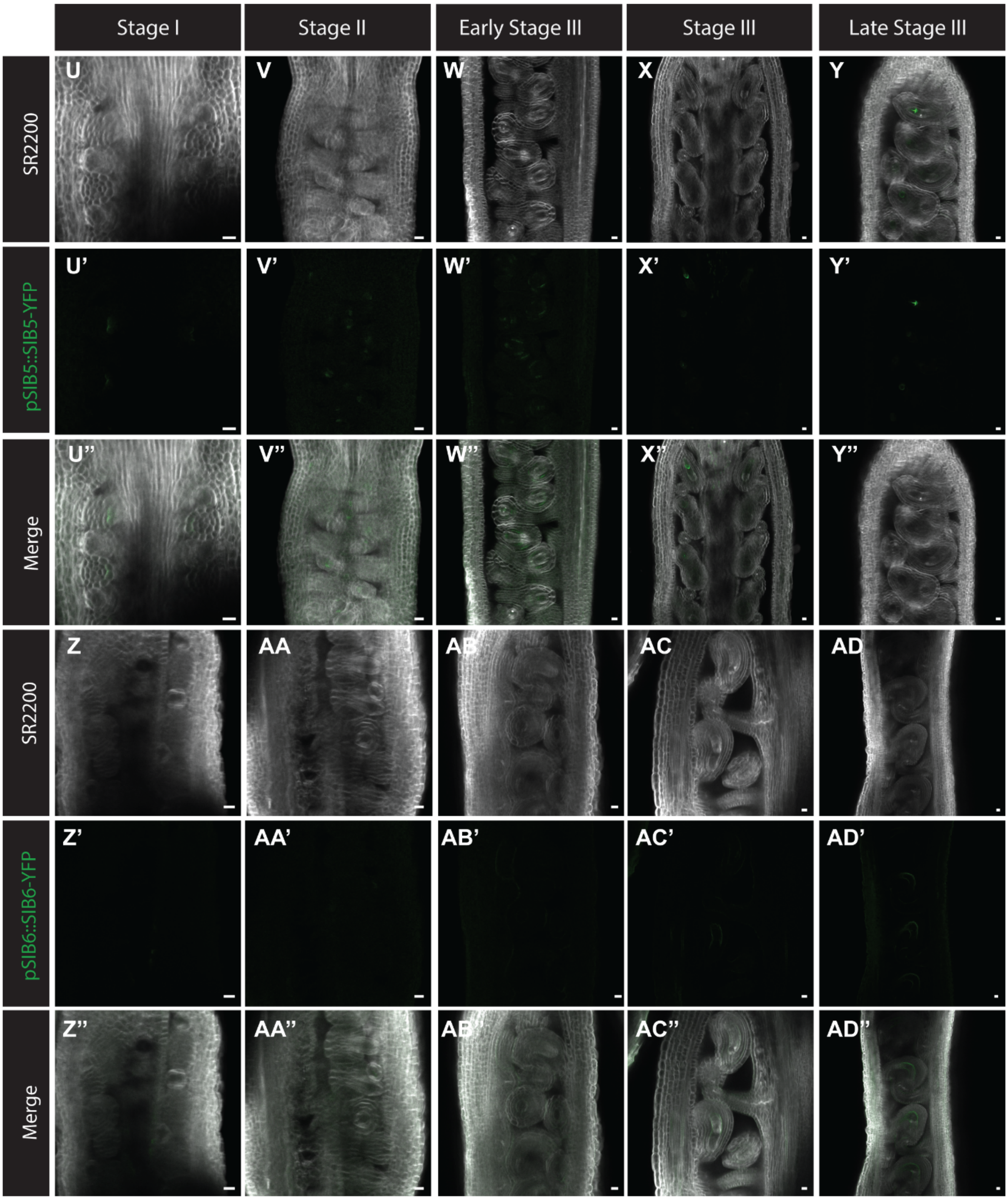

Stage I
Stage II
Early Stage III
Stage III
Late Stage III
SR2200
pSIB5::SIB5-YFP
Merge
SR2200
pSIB6::SIB6-YFP
Merge
U
V
W
X
Y
U'
V'
W'
X'
Y'
U''
V''
W''
X''
Y''
Z
AA
AB
AC
AD
Z'
AA'
AB'
AC'
AD'
Z''
AA''
AB''
AC''
AD''

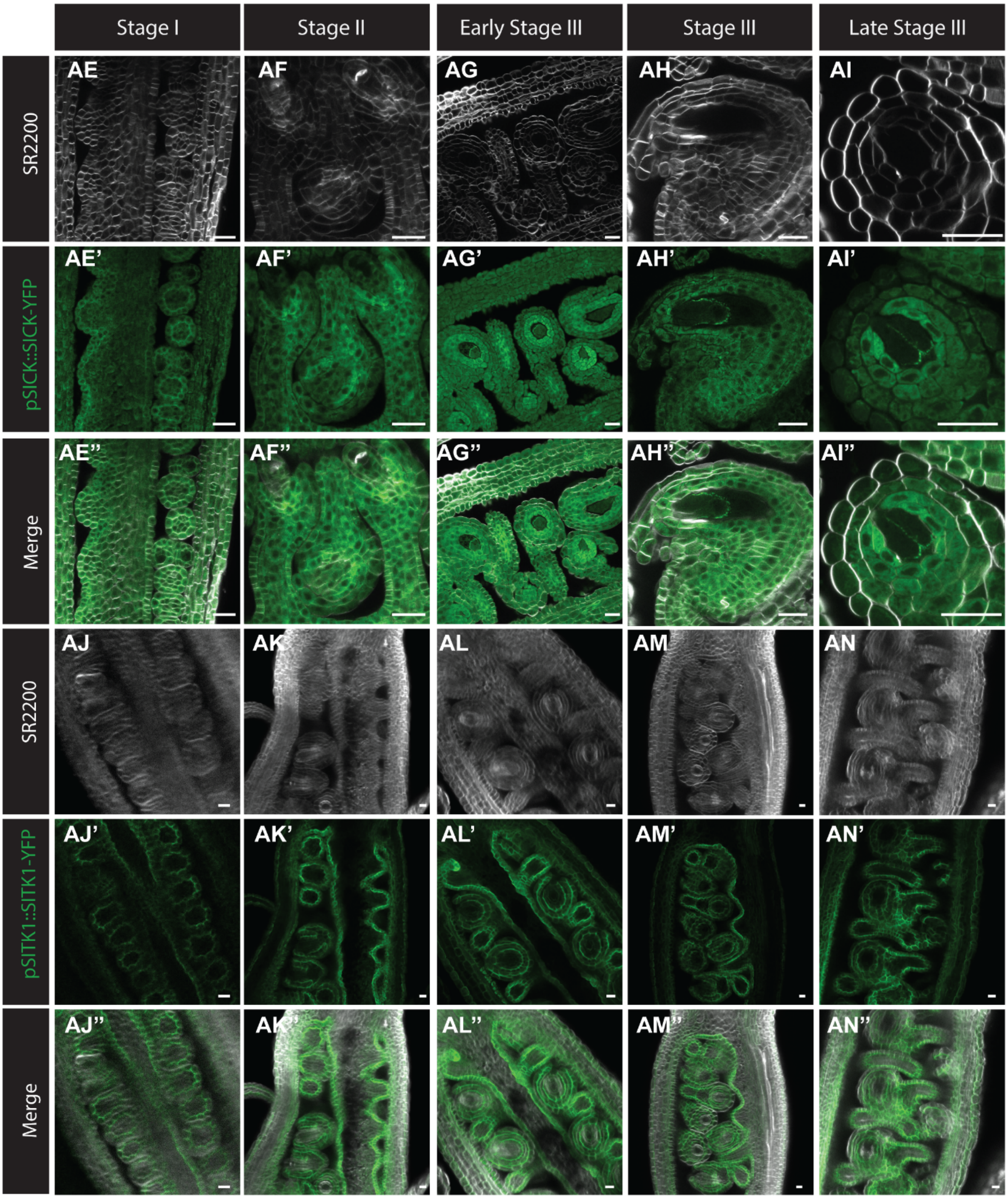

Stage I
Stage II
Early Stage III
Stage III
Late Stage III
SR2200
pSlCK::SlCK-YFP
Merge
SR2200
pSlTK1::SlTK1-YFP
Merge
AE
AF
AG
AH
AI
AE'
AF'
AG'
AH'
AI'
AE''
AF''
AG''
AH''
AI''
AJ
AK
AL
AM
AN
AJ'
AK'
AL'
AM'
AN'
AJ''
AK''
AL''
AM''
AN''

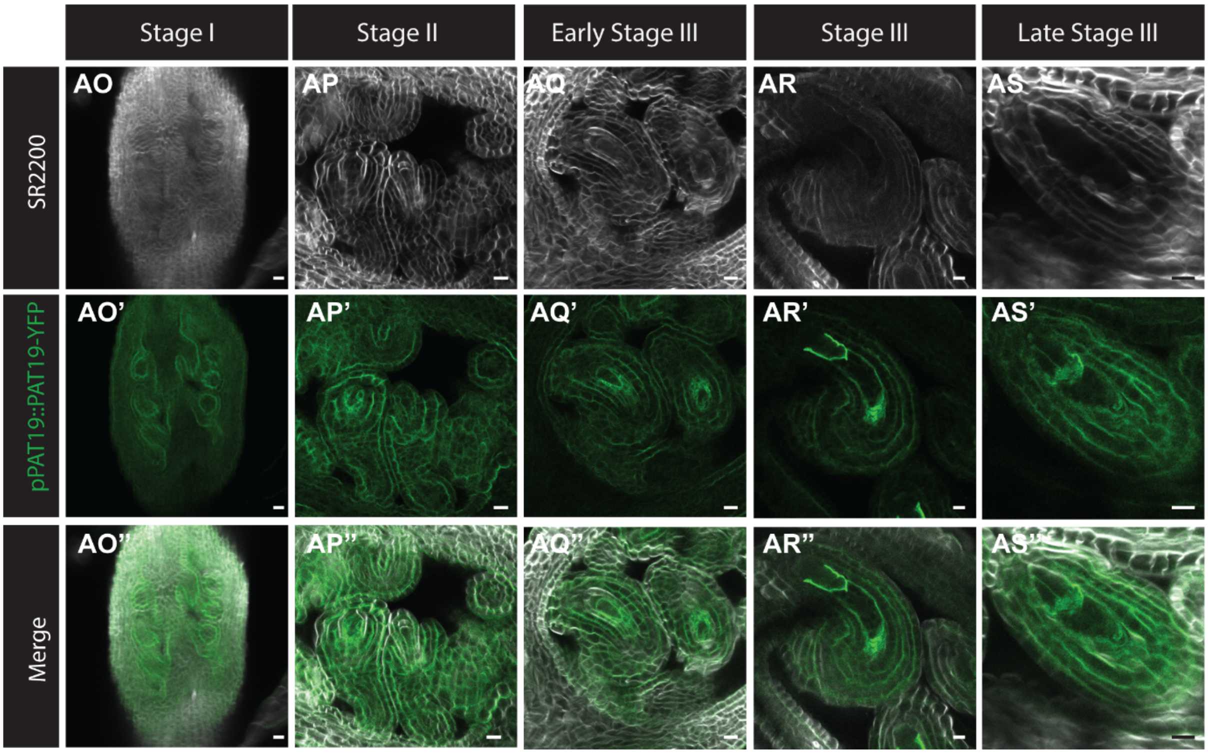


**Figure S3 Expression of SOK interacting proteins in the female gametophyte.** (A-AS) Confocal sections (seperated channels and merged images) of the developing ovule in stage I, stage II and III expressing pSIB1::gSIB1-YFP (A-E), pSIB2::gSIB2-YFP (F-J), pSIB3::gSIB3-TdT (K-O), pSIB4::gSIB4-YFP (P-T), pSIB5::gSIB5-YFP (U-Y), pSIB6::gSIB6-YFP (Z-AD), pSICK::gSICK-TdT (AE-AI), pSITK::gSITK-YFP (AJ-AN) and pPAT19::gPAT19-mSC (AO-AS). YFP, TdT, mSC, green; SR-2200, white. Images are displayed with brightness normalization and in some cases rotated or inverted to facilitate comparison of images. Tissues were fixed, cleared using Clearsee alpha (Kurihara et al., 2021) and stained with SR-2200 dye (Musielak et al., 2015) before imaging. Scale bars, 10 µm.

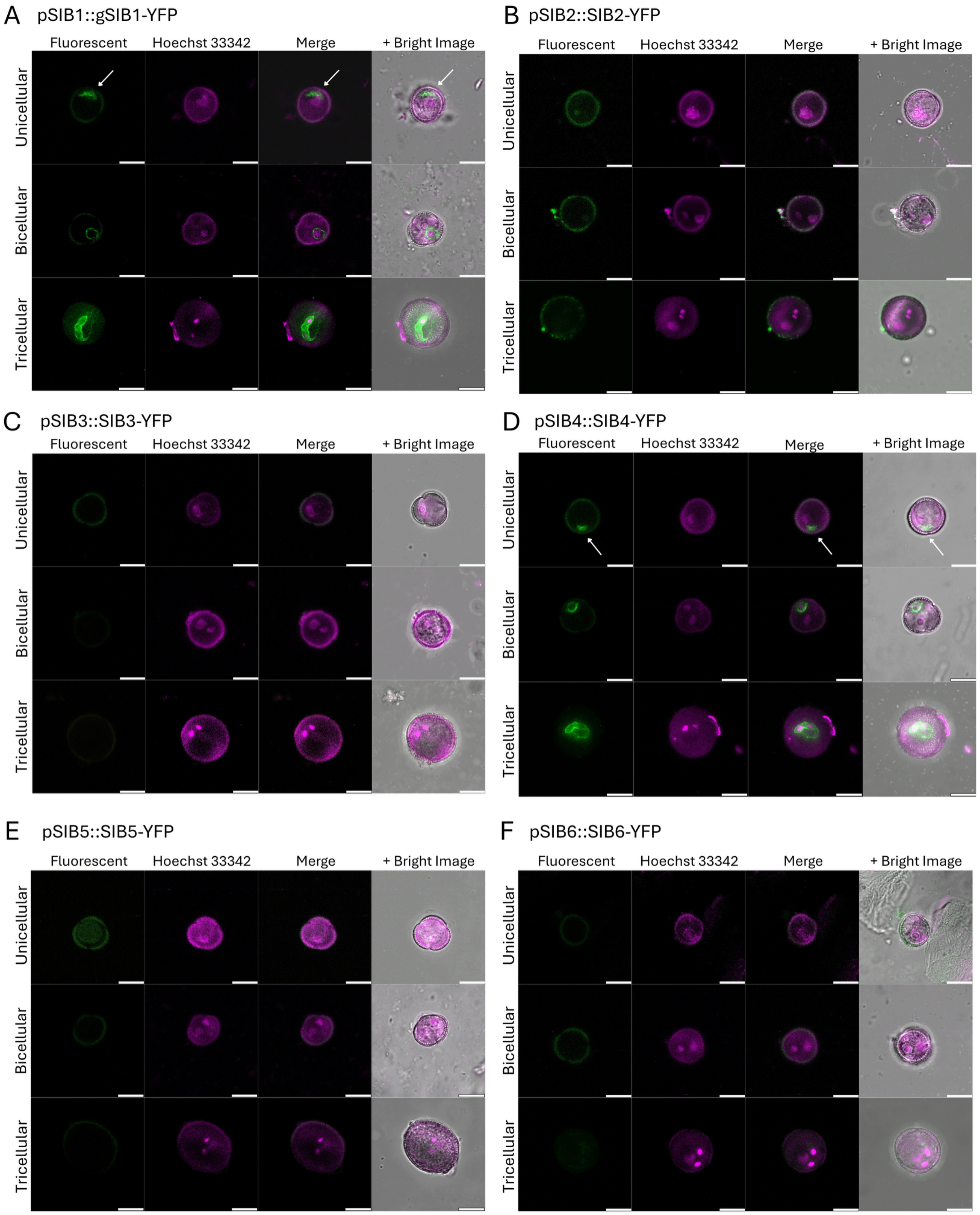


**Figure S4. Separated fluorescence channels for SIB reporters in the male gametophyte.** (A–F) Confocal sections of unicellular microspores, bicellular pollen and tricellular pollen expressing pSIB1::gSIB1-YFP (A), pSIB2::gSIB2-YFP (B), pSIB3::gSIB3-YFP (C), pSIB4::gSIB4-YFP (D), pSIB5::gSIB5-YFP (E) and pSIB6::gSIB6-YFP (F), shown as separated channels. Within each block, rows are the three developmental stages and columns are YFP fluorescence, Hoechst 33342, the two-channel merge, and the merge overlaid on the brightfield image. YFP, green; Hoechst 33342, magenta. Scale bars, 10 μm.

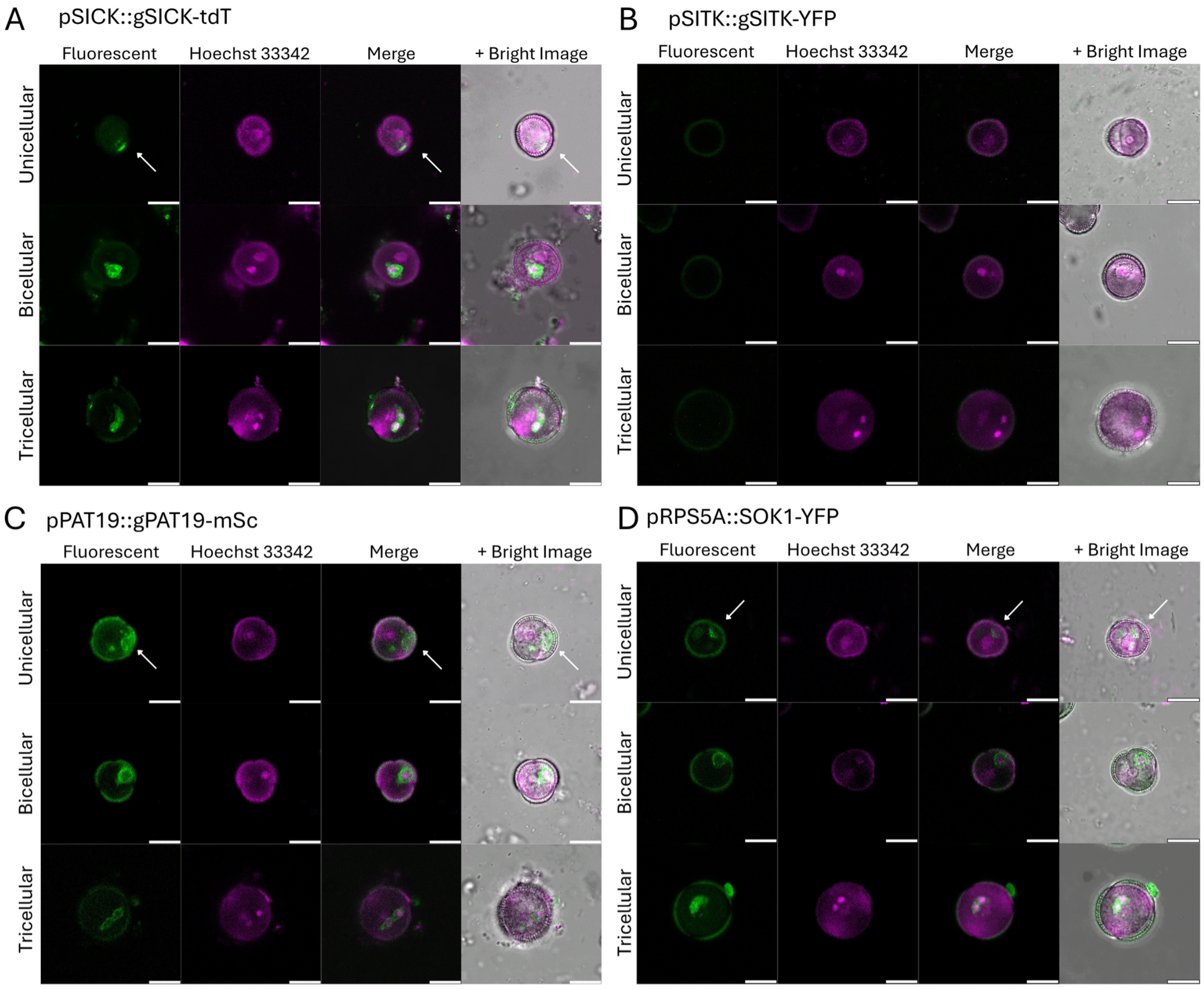


**Figure S5. Separated fluorescence channels for SITK1, PAT19 and SICK reporters and ubiquitously expressed SOK1 in the male gametophyte.** (A–D) Confocal sections of unicellular microspores, bicellular pollen and tricellular pollen expressing pSICK::gSICK-tdTomato (A), pSITK1::gSITK1-YFP (B), pPAT19::gPAT19-mScarlet (C), and pRPS5A::gSOK1-YFP (D), shown as separated channels. Within each block, rows are the three developmental stages and columns are reporter fluorescence, Hoechst 33342, the two-channel merge, and the merge overlaid on the brightfield image. The reporter is shown in green and Hoechst 33342 in magenta; note that PAT19-mScarlet and SICK-tdTomato are red-emitting fluorophores displayed in green for consistency with the YFP reporters. Scale bars, 10 µm.

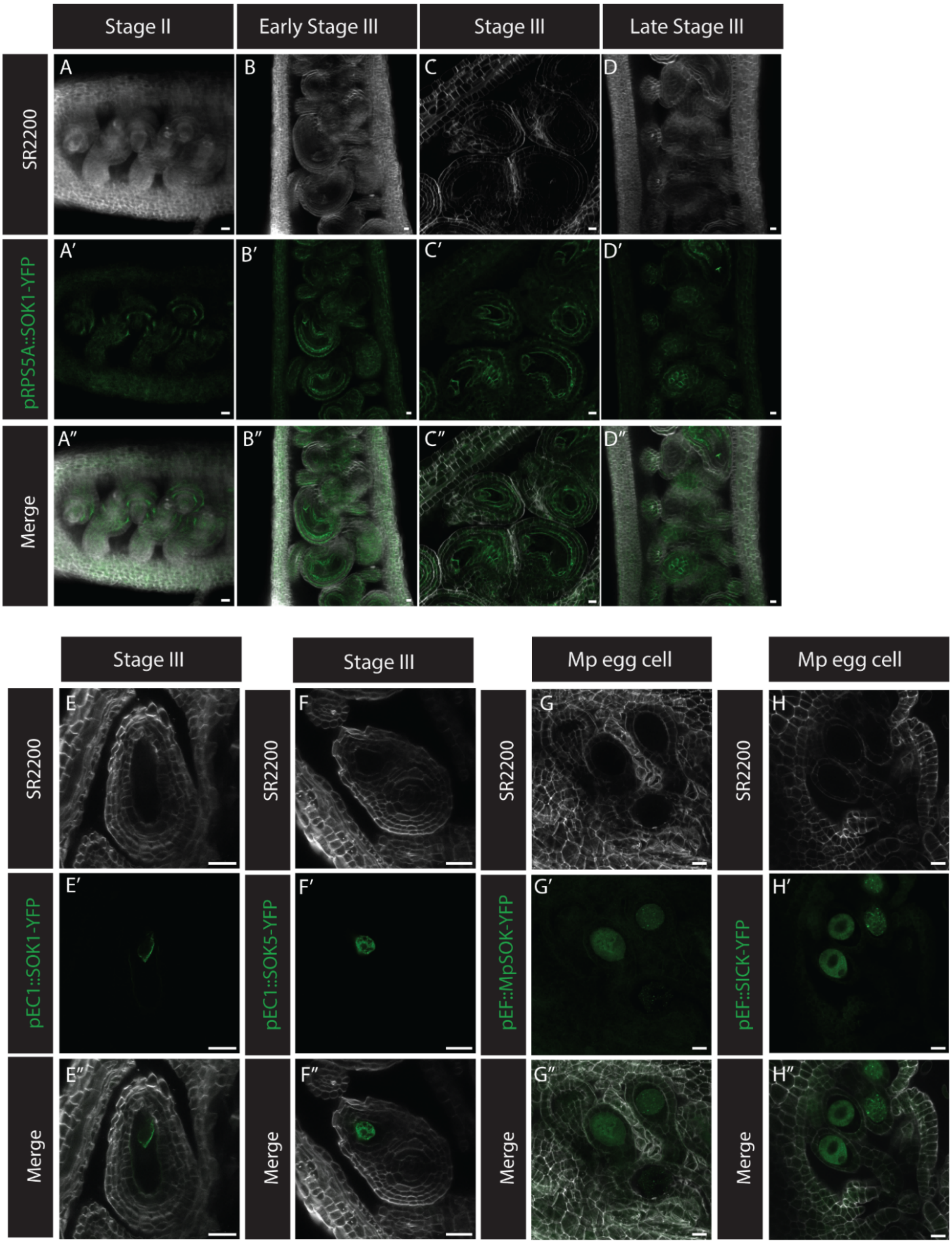


**Figure S6. Misexpression of SOK1 shows that the described polar complex is one system not multiple, furthermore egg cell polarity is not conserved during evolution .** (A-H) Confocal sections (seperated channels and merged images) of the developing ovule in stage II and III expressing pRPS5A::gSOK1-YFP. (A-D) Confocal sections of ovules in late stage III expressing pEC1::SOK1-YFP and pEC1::SOK5-YFP (E, F). Confocal sections of Marchantia egg cells, expressing EF::MpSOK-YFP and MpSICK::MpSICK-YFP (G, H). YFP, TdT, mSC, green; SR-2200, white; Hoechst 33342, magenta. Images are displayed with brightness normalization and in some cases rotated or inverted to facilitate comparison of images. Tissues were fixed, cleared using Clearsee alpha (Kurihara et al., 2021) and stained with SR-2200 dye (Musielak et al., 2015) before imaging. Scale bars, 10 μm.

**Table S1 Used primers**

| Primers for GoldenGate constucts | Sequence |
|---|---|
| SK166_GG_pSOK1_F1 | TATGAAGACACggagGAGTCGTTCCGTGGTGAATCAATG |
| SK167_GG_pSOK1_R5 | TATGAAGACACcattCTCTCTTTCTTTTTGTTTTGcTCTTCGATTTTATTCT |
| SK168_GG_pSOK1_R1 | GAATTGTTcTCTTCCATTTCTCTGATCG |
| SK169_GG_pSOK1_F2 | CGATCAGAGAAATGGAAGAgAACAATTC |
| SK170_GG_pSOK1_R2 | CATCTTCcTCTTCATACATAGACTTC |
| SK171_GG_pSOK1_F3 | GAAGTCTATGTATGAAGAgGAAGATG |
| SK172_GG_pSOK1_R3 | GGCTAATAGAAAGAAGAAGAgAAAAACATAG |
| SK173_GG_pSOK1_F4 | CTATGTTTTTcTCTTCTTCTTTCTATTAGCC |
| SK174_GG_pSOK1_R4 | CGACAAATCAGAAGAgAAAAAAAGGC |
| SK175_GG_pSOK1_F5 | GCCTTTTTTTcTCTTCTGATTTGTCG |
| SK207_GG_cSOK2_F1 | TATGAAGACACAATGATGGAAGCTGTAAGATGCAGAAGAG |
| SK208_GG_cSOK2_R1 | CAGTCTTGTTGACAATGTTCTTGC |
| SK209_GG_cSOK2_F2 | CATTGTCAACAAGACTGTCGTAGATGACGAG |
| SK210_GG_cSOK2_R2 | TATGAAGACACCGAACCTTTCTTGATTTGCTTCGATGATGACATTAAACAC |
| SK211_GG_cSOK4_F1 | TATGAAGACACAATGATGGCGTTGGTGAGTTCAAGAG |
| SK212_GG_cSOK4_R1 | CTTTGAGATACAGACCGTTGTGAG |
| SK213_GG_cSOK4_F2 | CGGTCTGTATCTCAAAGATGTAATCAATAG |
| SK214_GG_cSOK4_R2 | TATGAAGACACCGAACCTGGTTGTCCAACCACAGAATTTGG |
| SK215_GG_CSOK3_F | TATGAAGACACAATGATGGAAGCGAGGATGAAGAAGTATAGC |
| SK216_GG_CSOK3_R | TATGAAGACACCGAACCAGGCTCTTCTGAAGGCACTT |
| SK217_GG_cSOK5_F | TATGAAGACACAATGATGAGTTCAAGAGTGTTCAGAGCAAC |
| SK218_GG_cSOK5_R | TATGAAGACACCGAACCCTGCTGCCCACCATTGTTC |
| **Primers for ligation-independent cloning contructs** | **Sequence** |
| SIB4pro_F | AATTCTAGTTGGAATGGGTTACCAAAAGCCCAAAACCGTC |
| SIB4pro_R | CCAAGTCTCATAAACGCCATTGTGGAAGAAAGTCTTGAGTTGGTG |
| SIB4cds_F | ACTCAAGACTTTCTTCCACAATGGCGTTTATGAGACTTGGCTC |
| SIB4cds_R | ACAGCTCCTCGCCCTTACTAGTCATAGAACCTCCCCCACCAGAACCCCCACCAATTGCTGGTGTTGCTTGATCC |
| SIB5pro_F | AATTCTAGTTGGAATGGGTTAGGTGTTAGGGGAGAGAGAAGA |

| | |
|---|---|
| SIB5pro_R | CCCAGTTTCAAGCTAGCCATCTTTGTTCTTTCACTTCAACTCTTTGGTT |
| SIB5cds_F | GTTGAAGTGAAAGAACAAAGATGGCTAGCTTGAAACTGGGTTC |
| SIB5cds_R | ACAGCTCCTCGCCCTTACTAGTCATAGAACCTCCCCCACCAGAACCCCCACCATTTTTGGTTTCAGGAATCACTAGAGC |
| SIB6pro_F | AATTCTAGTTGGAATGGGTTTGGTGGCTGCATTGGAGAAT |
| SIB6pro_R | TGCTTCTTGGAGTTAGCCATTGGAGAAAATCACAGAAAAAAACAAAGAAAG |
| SIB6cds1_F | GTTTTTTTCTGTGATTTTCTCCAATGGCTAACTCCAAGAAGCAAGC |
| SIB6cds1_R | TCCAATATCCATGAATGAGTCTTGAAATATATATCGACGGCTCTGTATAG |
| SIB6cds2_F | CCGTCGATATATATTTCAAGACTCATTCATGGATATTGGAGGAAGAGA |
| SIB6cds2_R | ACAGCTCCTCGCCCTTACTAGTCATAGAACCTCCCCCACCAGAACCCCCACCACTAAGAAAACGTCGTGATCTTCTATCG |
| **Primers for Gateway cloning constructs** | **Sequence** |
| CA1467 - AtSOK1 att fw | GGGGACAAGTTTGTACAAAAAAGCAGGCTTCATGGAAAGTAATGGTGGAGGAGGAGAAG |
| CA1468 - AtSOK1 att rv | GGGGACCACTTTGTACAAGAAAGCTGGGTCCTCTTTGAGAGTAGTCGTCAATACACGTTG |
| CA1475 - AtSOK5 att fw | GGGGACAAGTTTGTACAAAAAAGCAGGCTTCATGAGTTCAAGAGTGTTCAGAGCAACACCAG |
| CA1476 - AtSOK5 att rv | GGGGACCACTTTGTACAAGAAAGCTGGGTCCTGCTGCCCACCATTGTTCCTTTTAG |
| CA863 - EC1 fw | TAGCAAGCTTTAGATGACGGAATTAGCATATCTCATGCAC |
| CA864 - EC1 rv | AGCTAAGCTTTTCTCAACAGATTGATAAGGTCGAAAGAAG |
